# Molecular phenotypes segregate missense mutations in SLC13A5 Epilepsy

**DOI:** 10.1101/2024.05.23.594637

**Authors:** Valeria Jaramillo-Martinez, Souad R. Sennoune, Elena B. Tikhonova, Andrey L. Karamyshev, Vadivel Ganapathy, Ina L. Urbatsch

## Abstract

The sodium-coupled citrate transporter (NaCT, SLC13A5) mediates citrate uptake across the plasma membrane via an inward Na^+^ gradient. Mutations in SLC13A5 cause early infantile epileptic encephalopathy type-25 (EIEE25, SLC13A5 Epilepsy) due to impaired citrate uptake in neurons. Despite clinical identification of disease-causing mutations, underlying mechanisms and cures remain elusive. We mechanistically classify the molecular phenotypes of six mutations. C50R, T142M, and T227M exhibit impaired citrate transport despite normal expression at the cell surface. G219R, S427L, and L488P are hampered by low protein expression, ER retention, and reduced transport. Mutants’ mRNA levels resemble wildtype, suggesting post-translational defects. Class II mutations display immature core-glycosylation and shortened half-lives, indicating protein folding defects. These experiments provide a comprehensive understanding of the mutation’s defects in SLC13A5 Epilepsy at the biochemical and molecular level and shed light into the trafficking pathway(s) of NaCT. The two classes of mutations will require fundamentally different treatment approaches to either restore transport function, or enable correction of protein folding defects.

**Summary:** Loss-of-function mutations in the SLC13A5 causes SLC13A5-Epilepsy, a devastating disease characterized by neonatal epilepsy. Currently no cure is available. We clarify the molecular-level defects to guide future developments for phenotype-specific treatment of disease-causing mutations.

## Introduction

### Importance of citrate to brain function

Citrate is a key intermediate in the tricarboxylic acid (TCA) cycle that connects carbohydrate and fat metabolism (Tong and Rouault, 2007). This metabolite is found not only in mitochondria and cytoplasm within select cells but also in blood plasma (100–200 µM) (Fraenkl et al., 2011; Hamm, 1990). Citrate in plasma is transported into cells by the Na^+^-coupled citrate transporter (NaCT) encoded by the gene *SLC13A5*. NaCT is expressed in the brain, liver, bones and teeth, and testes, where it delivers citrate across the cell membrane into the cytoplasm. In the brain where NaCT is expressed primarily in astrocytes and to some extent in neurons depending on the species examined, citrate serves as an energy source as well as a precursor for the synthesis of the neurotransmitters acetylcholine, GABA, and glutamate (Yodoya et al., 2006). In the liver, citrate is involved in energy production and cholesterol/fatty acid synthesis (Bhutia et al., 2017; Gopal et al., 2007; Higuchi et al., 2020; Willmes et al., 2018). In bones and teeth, citrate chelates calcium and helps in the mineralization of these tissues (Irizarry et al., 2017). In the testes, citrate helps with the energy needs of the sperm. Until recently it was assumed that citrate found in the cytoplasm originates solely from mitochondria via transport mediated by the mitochondrial citrate transporter SLC25A1 (also known as CIC or citrate carrier). The identification of the plasma membrane citrate transporter NaCT has brought focus to extracellular citrate as an important source of cytoplasmic citrate in select cells. Importantly, recent studies have shown differential compartmentalization and biological functions of cytoplasmic citrate pools in brain depending on whether they originate via SLC25A1/CIC or SLC13A5/NaCT (Fernandez-Fuente et al., 2023). These recent findings underscore the importance of NaCT to brain function. Loss-of-function mutations in NaCT cause early infantile epileptic encephalopathy type 25 (EIEE25, SLC13A5 Epilepsy), an autosomal recessive disease, in humans (Bhutia et al., 2017; Hardies et al., 2015; Jaramillo-Martinez et al., 2021c; Klotz et al., 2016; Kopel et al., 2021; Selch et al., 2018; Spelbrink et al., 2023; Thevenon et al., 2014; Willmes et al., 2018). The disease is characterized by neonatal epilepsy, delayed brain development and language skills, defective bone mineralization, and abnormal tooth development and enamalization.

### Clinical phenotypes of disease-causing mutations in NaCT

SLC13A5 Epilepsy is a rare disease with about 145 patients registered in the database (TESS Research Foundation), carrying 35 sequence-confirmed missense mutations that cause disease. The clinical phenotypes vary between patients, and detailed descriptions are not fully available. Among these mutations, G219R is the most frequent found in 31 patients who carry the mutation on at least one allele, of which 22 have been studied (Klotz et al., 2016; Matricardi et al., 2020; Thevenon et al., 2014; Weeke et al., 2017; Yang et al., 2020), see **Table 1**. Patients carrying homozygous alleles (three patients) showed generalized tonic-clonic seizures once a week, and most of them died at an early age. Patients with compound heterozygosity (19 patients) presented seizures over a wide range of frequencies, from 50 to 100 times a day to just once a year (Klotz et al., 2016; Matricardi et al., 2020; Thevenon et al., 2014; Weeke et al., 2017; Yang et al., 2020). The data suggest that the heterozygous mutant variant may determine severity of the disease. Other common mutations include S427L, T142M, T227M, C50R, and L488P as detailed in **Table 1**. S427L is found in seven patients, three homozygous and four compound heterozygous; all patients present with daily seizures (Hardies et al., 2015; Matricardi et al., 2020; Weeke et al., 2017). T142M is found as a homozygous mutation in two patients, and more frequently as a compound heterozygous mutation (six patients). The phenotypes associated with this mutation vary from no seizures to everyday seizures with no clear genetic link to the heterozygous variant (Matricardi et al., 2020; Weeke et al., 2017; Yang et al., 2020). T227M is found in six heterozygous patients and two homozygous patients. The phenotypes, similar to T142M, vary from no seizures to everyday seizures (Klotz et al., 2016; Matricardi et al., 2020; Thevenon et al., 2014; Weeke et al., 2017; Yang et al., 2020). Homozygous C50R mutation is found in two patients. One of these patients exhibited seizures when fever was present or when antiepileptic medication was not administered whereas the other patient only displayed seizures until five years of age (Yang et al., 2020). Patients with the L488P mutation are homozygous from the same family and present with weekly to monthly seizures (Thevenon et al., 2014). Despite the clinical identification of many of these NaCT mutations in patients, their disease-causing mechanisms are not clear.

**Table 1.**
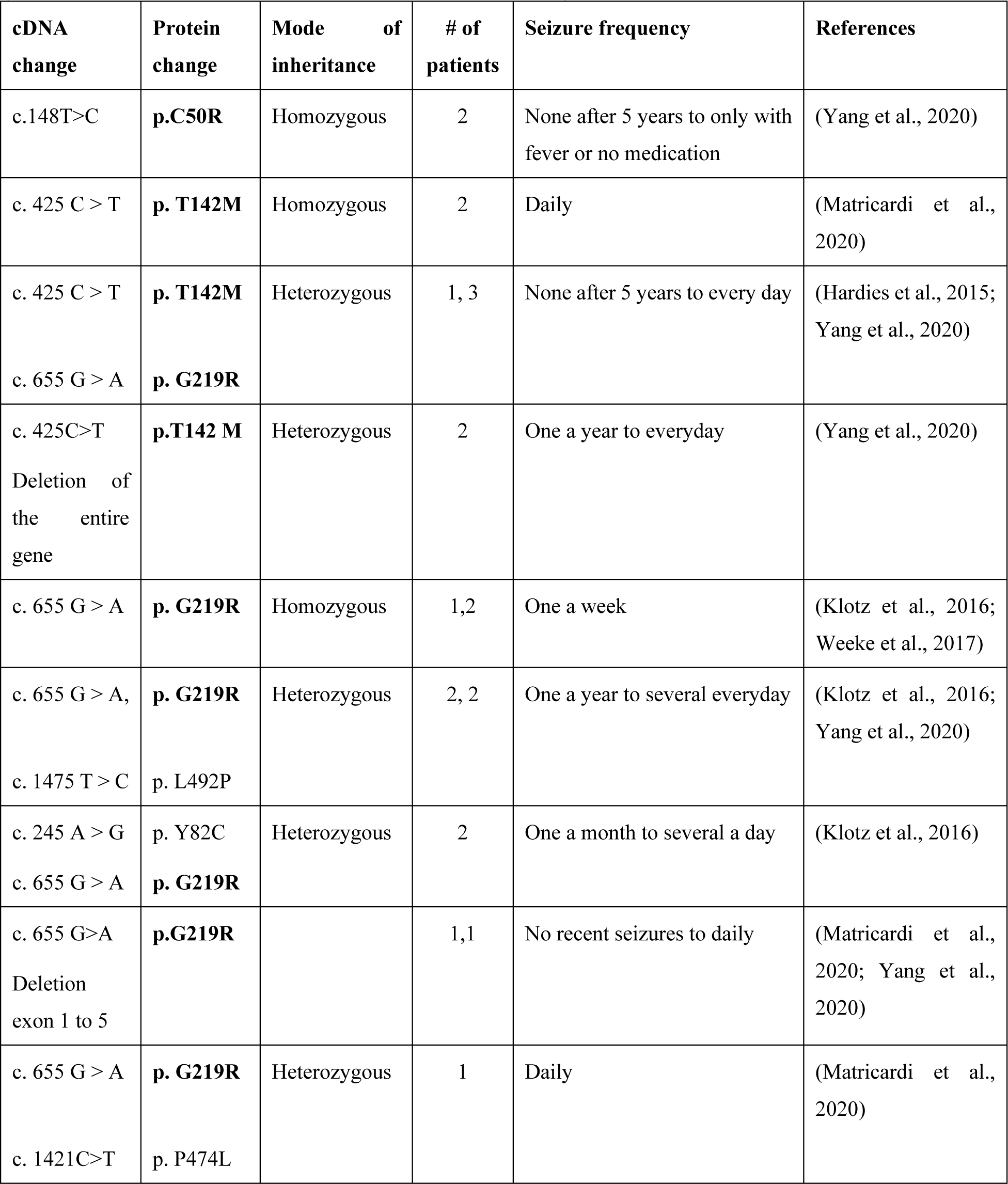
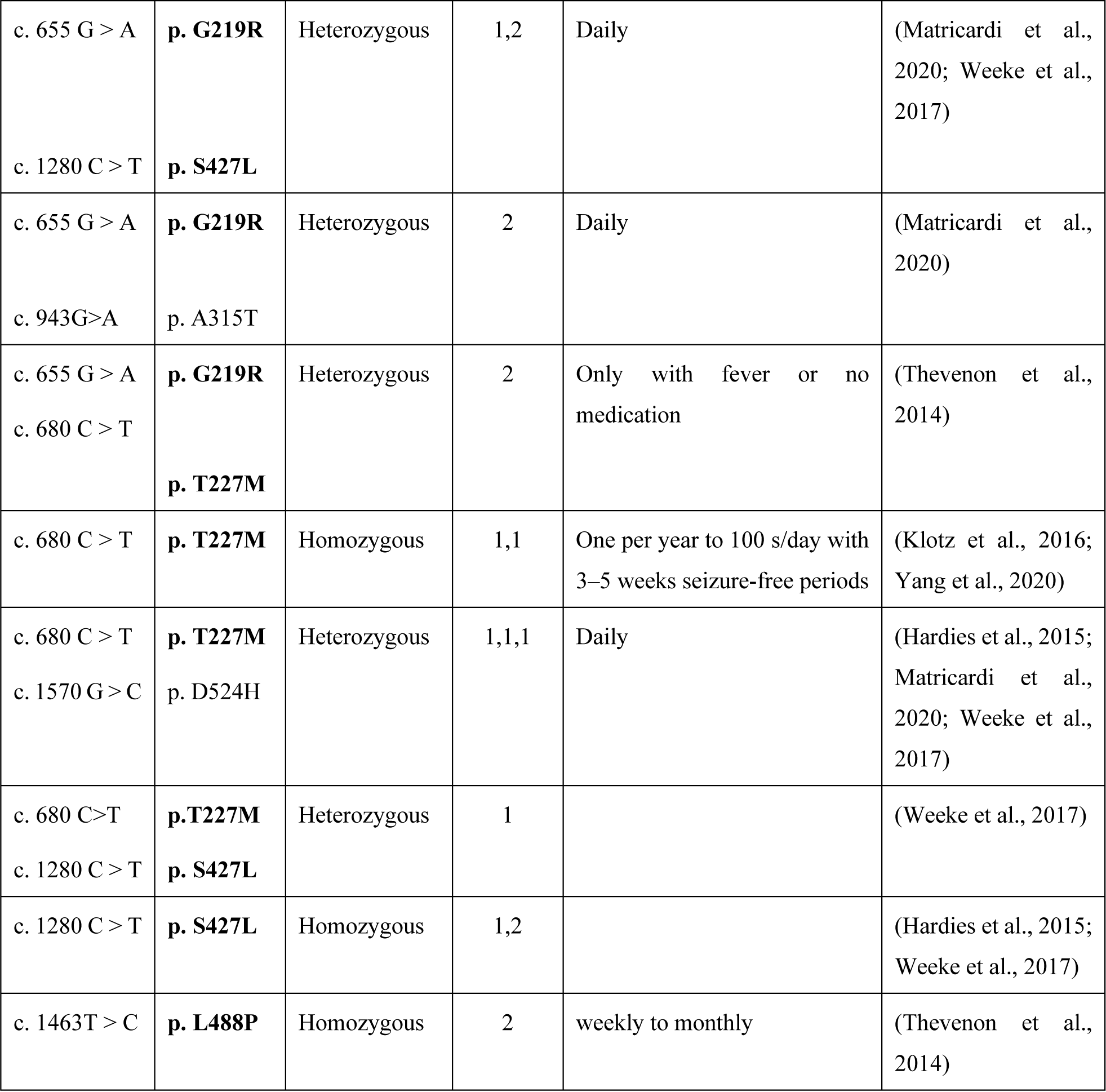
Clinical phenotypes and frequency of six of the most common missense mutations (C50R, T142M, G219R, T227M, S427L, and L488P).

To date, no cure is available for this devastating disease. To develop effective personalized treatments, we need to fully understand, at the molecular level, the underlying defect of missense mutations. This will require detailed analyses of the consequences of these mutations in terms of cell biology and biochemical functions of NaCT. Only a few studies have scantly reported on the protein expression and transport malfunctions of disease causing SLC13A5 Epilepsy mutations, yet with contradictory results (Hardies et al., 2015; Klotz et al., 2016; Selch et al., 2018). In part, this is due to the lack of highly sensitive NaCT-specific antibodies that can unequivocally recognize low levels of the human protein by immunofluorescence and Western blots. In particular, the expression profiles of the mutations G219R, L488P and T227M as well as the molecular sizes of the full-length, mature proteins are controversal. Furthermore, none of the prior reports considered the glycosylation status of the mutant proteins or lack thereoff, that may serve as an important clue to traffiquing defects. In the present study, we focused on the six most common mutations C50R, T142M, G219R, T227M, S427L, and L488P (**Table 1**). We clarify and refine the cell biology and biochemical phenotypes by probing their transport functions and expression profiles compared to wildtype NaCT. We go beyond existing studies and investigated sodium-coupled and lithium-stimulated transport activities in transiently transfected and clonal cell lines, and we further analyze surface expression, cellular localization patters using multiple marker proteins, as well as mRNA synthesis, protein life times, and protein trafficking and degradation pathways.

## Results

### Location of disease-causing mutations in the NaCT cryo-EM structure

NaCT is a homodimer with each subunit comprised of a transporter domain and scaffold domain (**Fig. 1A**). The scaffold domain functions as an anchor for the transporter in the membrane, while the transport domain is responsible for binding and moving substrates across the membrane. The relative location of the six major missense mutations (C50R, T142M, G219R, T227M, S427L, and L488P) in the primary amino acid sequence, and the single nucleotide/codon changes (red) are displayed in **Fig. 1B**. These mutations are distributed throughout the protein, with T142M, G219R, T227M, and L488P localized within the transporter domain (yellow in **Fig. 1B**), and C50R and S427L situated in the scaffold domain (blue). Notably, mutations T142M and T227M (green sticks) localize close to the two sodium-(purple balls Na1 and Na2) and the citrate-binding sites (orange sticks) and are within a binding distance of the citrate in the cryo-EM structure of human NaCT PDB: 7JSK (Sauer et al., 2021a) (**Fig. 2A**). While G219R is in proximity to the Na1 site, it is somewhat farther away (12 Å) to citrate. L488P and S437L in the C-terminal transport domain are even further away from citrate (22.8 Å and 30.5 Å, respectively) and are likely not involved in the binding of Na^+^ and/or citrate (**Fig. 4A**). C50R located in the scaffold domain (blue) appears to reside at the interface between the two domains (Jaramillo-Martinez et al., 2021a; Jaramillo-Martinez et al., 2021c). Although the recently determined high-resolution structure of NaCT gave a clue of how point mutations might directly pertube the transport function if located near the substrate binding site, it is prudent to dissect the mutant mechanisms in functional assays, especially for those that are further away from the citrate transport pathway.

**Figure 1.**
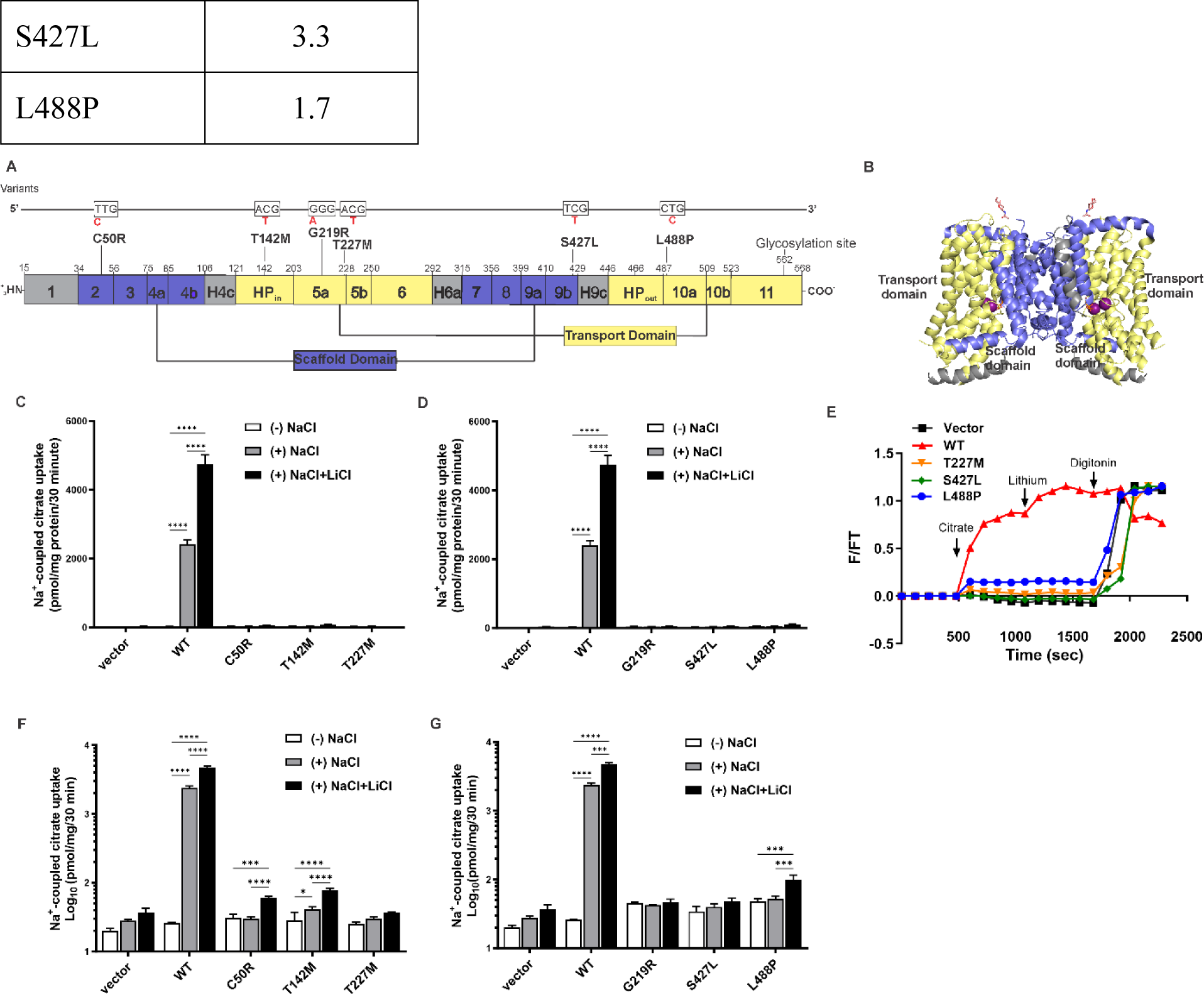
Differential transport activity of WT and mutant NaCTs. (**A**) 3D structure of the NaCT dimer (PDB: 7JSK,(Sauer et al., 2021a)); Na^+^ ions are shown as magenta balls, and citrate is in orange sticks. (**B**) Location of six most frequent missense mutations in the primary sequence of NaCT with the scaffold domain colored in blue, the transport domain in yellow. Transmembrane helices H4c, H6a, and H9c (Elsby et al.) bridge the transport domain to the scaffold domain. WT codons are shown above in black with nucleotide substitutions indicated in red. color coded as in (**A**). (**C**) and (**D**) Na^+^-coupled uptake of [^14^C]-citrate in HEK293 cells transiently transfected with NaCT mutants as described in Methods. (**E**) Fluorescence time course of citrate uptake in HEK293 cells co-transfected with Citron and either empty vector, WT, or mutant variants. Relative fluorescence changes were measured in NaCl buffer before and after consecutive additions of 10 mM citrate and 10 mM Li^+^ (arrows), followed by 0.5% digitonin to reveal maximally achievable total fluorescence(Knauf et al.). (**F**) and (**G**) Log-scale of (**C**) and (**D**) to reflect the very low activity of mutant variants. Each point represents the mean ± SEM of 3-5 experiments *P ≤ 0.05, **P ≤ 0.01, and ***P ≤ 0.001. In (E) error bars were small and were omitted for clarity.

**Figure 2.**
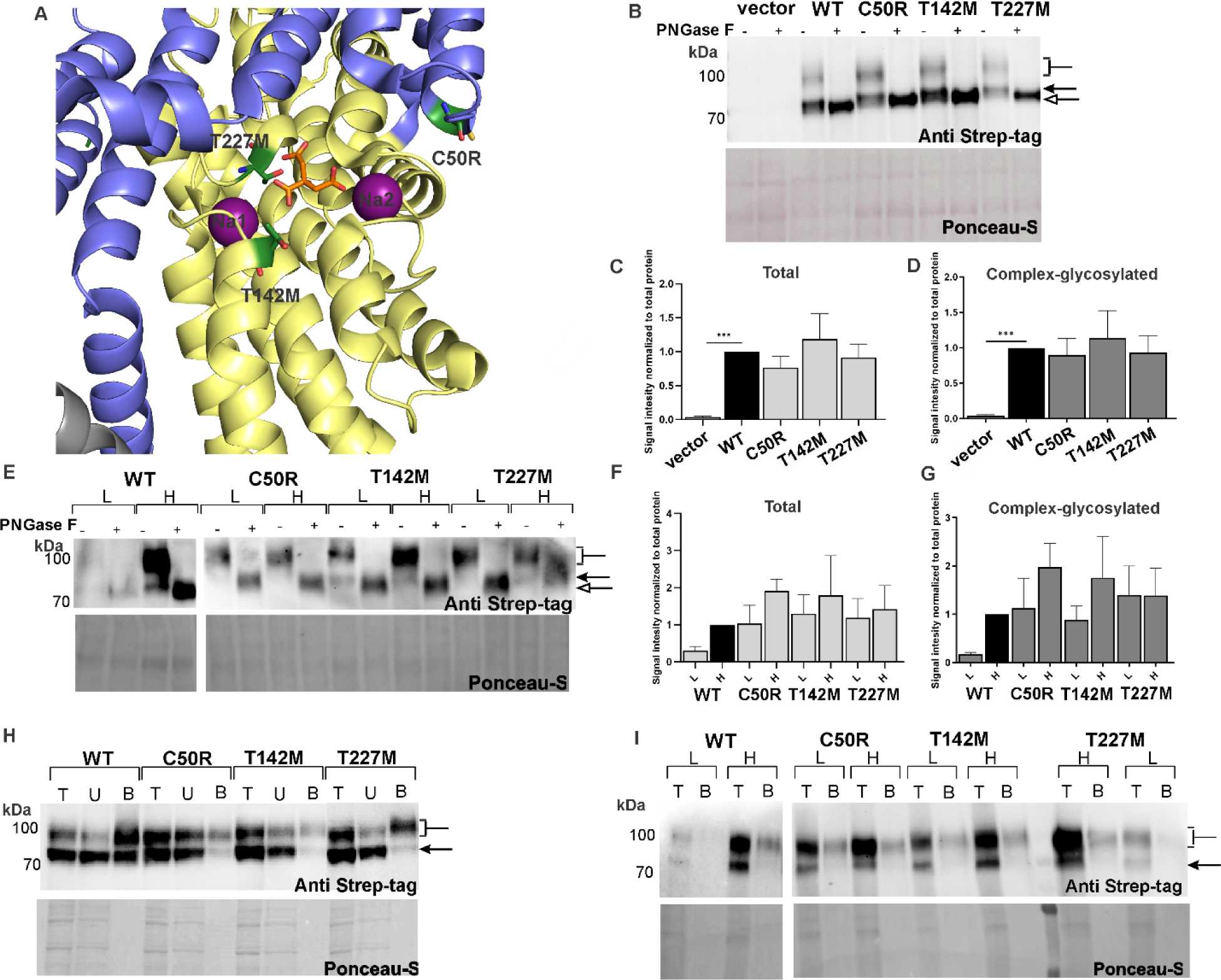
Glycosylation status and expression profiles of Class I mutants. (**A**) Location of C50R, T142M, and T227M in the cryo-EM structure (PDB: 7JSK) (Sauer et al., 2021a), scaffold and transport domains are colored as in Figure 1. (**B**) Whole-cell lysates from transiently transfected cells were treated without or with PNGase F, resolved by SDS-PAGE, and Western blots developed with the NaCT-specific anti-Strep-tag antibody. The sizes of non-glycosylated (empty arrow), core-glycosylated (arrow), and complex-glycosylated (brackets) NaCT protein bands are indicated. Quantification of (**C**) total NaCT Western blot signals and of (**D**) the complex-glycosylated bands from three independent experiments as decribed in Methods. (**E**) Western blot analysis of whole-cell lysates from clonal cell lines expressing lower (L) and higher (H) level of NaCT variants. Quantification of (**F**) total NaCT signals and the (**G**) complex-glycosylated protein bands. Total proteins loaded per lanes were stained with Ponceau S and used as loading controls. Cell-surface biotinylated proteins expressed in (**H**) transiently transfected and (**I**) clonal cells. Total protein lysates (T), unbound proteins (U) that washed off the NeutrAvidin resin, and bound biotinylated NaCT (B) were detected with the NaCT-specific anti-Strep-tag antibody. Each column represents the mean ± SEM, n= 3. *P ≤ 0.05, **P ≤ 0.01, and ***P ≤ 0.001. The relative levels were quantified by Image J, n=3.

### SLC13A5 Epilepsy causing mutations suffer severe loss of sodium-coupled citrate transport

We measured the citrate transport activity of the six major SLC13A5 disease-causing mutations. To quantitatively assess loss of function, we monitored [^14^C]-citrate uptake in HEK293 cells that transiently expressed either WT NaCT or mutant proteins. In the absence of Na^+^, very low level of citrate uptake was detected in all samples, similar to control cells transfected with the “empty” pcDNA3.1 vector (**Fig. 1C**, **D**, note replot of the same data in log scale in **Fig. 1F**, **G**). In the presence of Na^+^ (140 mM), robust citrate uptake into cells with WT NaCT was observed accumulating more than 2,000 pmol of citrate per mg of cellular protein. In contrast, citrate accumulation in cells expressing mutants C50R, T142M or T227M (**Fig. 1C**, **F**) or in G219R, S427L or L488P NaCTs (**Fig. 1D**, **G**) was very low, and was close to the detection limit compared to ‘no Na” samples and to samples transfected with pcDNA vector. To stimulate citrate transport, LiCl was added to NaCl-containing uptake buffers. We previously demonstrated that adding Li^+^ will specifically stimulate citrate transport mediated by human NaCT in the presence of sodium (Gopal et al., 2015; Inoue et al., 2003). At a concentration of 10 mM, Li^+^ stimulated uptake of Na^+^-coupled citrate by about two-fold in cells expressing WT NaCT. Interestingly, some low level of Li-stimulated citrate uptake was detectable for C50R, T142M and L488P mutants. The data suggest that Li may be incorporated in treatment strategies for these mutants.

Similar results were obtained using a green fluorescence protein variant “Citron” that detects accumulation of citrate in intact cells. For this HEK293 cells were co-transfected with plasmid coding for Citron and either pcDNA3.1 ‘empty’ vector, or WT NaCT or mutant variants. We demonstrate a rapid increase of fluorescence within minutes of adding citrate to cells only if they express functional WT NaCT (red in **Fig. 1E**). Vector control cells (black in **Fig. 1E**) or the mutants T227M and S427L showed very little change in fluorescence. Interestingly, the mutant L488P appeared to display some low-level activity in repeat experiments (blue in **Fig. 1E**).

### Expression of glycosylated mutant protein serves as first tier evaluation

To determine the primary cause of loss of transport function, protein expression was analyzed by Western blotting to assess total protein content in cell lysates with respect to glycosylation status. This gives an excellent first tier evaluation of whether a mutant potein is glycosylated, which indicates that synthesis/cotranlational folding in the ER is normal, and is followed by trafficking through the Golgi network where glycosylation maturates on the way to the plasma membrane.

Robust expression, similar to WT NaCT, was observed for mutants C50R, T142M, and T227M (**Fig. 2**). Initially, cell lysates of transiently transfected HEK293 cells were harvested 48 h post-transfection, and analyzed by Western bloting with the monoclonal anti-Strep-tag antibody that recognizes an epitope engineered to the N-terminus of NaCT (see Methods). Cells transfected with WT NaCT showed two bands that were absent in cells transfected with the “empty” pcDNA3.1 vector (**Fig. 2B**). If the same lysates were treated with PNGase F, which cleaves the innermost N-acetylglucosamine from asparagine, the top protein band collapsed to a lower size than the core glycosylated (named non-glycosylated, empty arrow). NaCT contains a predicted N-glycosylation site at N562. PNGase F treatment demonstrated that both bands observed without treatment are glycosylated proteins, and suggested that the upper band is complex and the lower is core glycosylated. Similarly, the three mutants C50R, T142M, and T227M all showed robust expression of two bands in cell lysates (**Fig. 2B**, **C**, and **D**). Quantification of the Western blot signals of the upper and lower NaCT protein bands suggested that the total signal in both bands (**Fig. 2C**) as well as the amount of complex glycosylated proteins (**Fig. 2D**) were comparable between mutants and wildtype. The data infer that the translated WT and mutant NaCT proteins attained complex glycosylation and matured while trafficking through the Golgi apparatus.

Transiently transfected cells are known to harbor a heterogeneous mixture of low, high, and non-expressing cells. To verify that phenotypes persist independently of expression levels, we generated cell lines stably expressing NaCT at lower and higher levels (**Supplemental Fig. S1**). Notably, in clonal cell lines, complex glycosylated protein (upper bands ∼90 kDa in **Fig. 2E** and **Supplemental Fig. S1**) were much more prominent than core glycosylated protein bands in all cases, whether expression levels were low or high, indicating the amount of protein synthesis was not limiting escape of the protein from the ER and trafficking through the Golgi apparatus. Notably, higher expression level of the WT NaCT protein in clonal cell lines correlated with a 2-fold increase of transport activity (**Fig. 3A and C**). However, higher-level expression of mutants T142M or T227M in clonal cell lines resultated in only marginally increased citrate transport.

**Figure 3.**
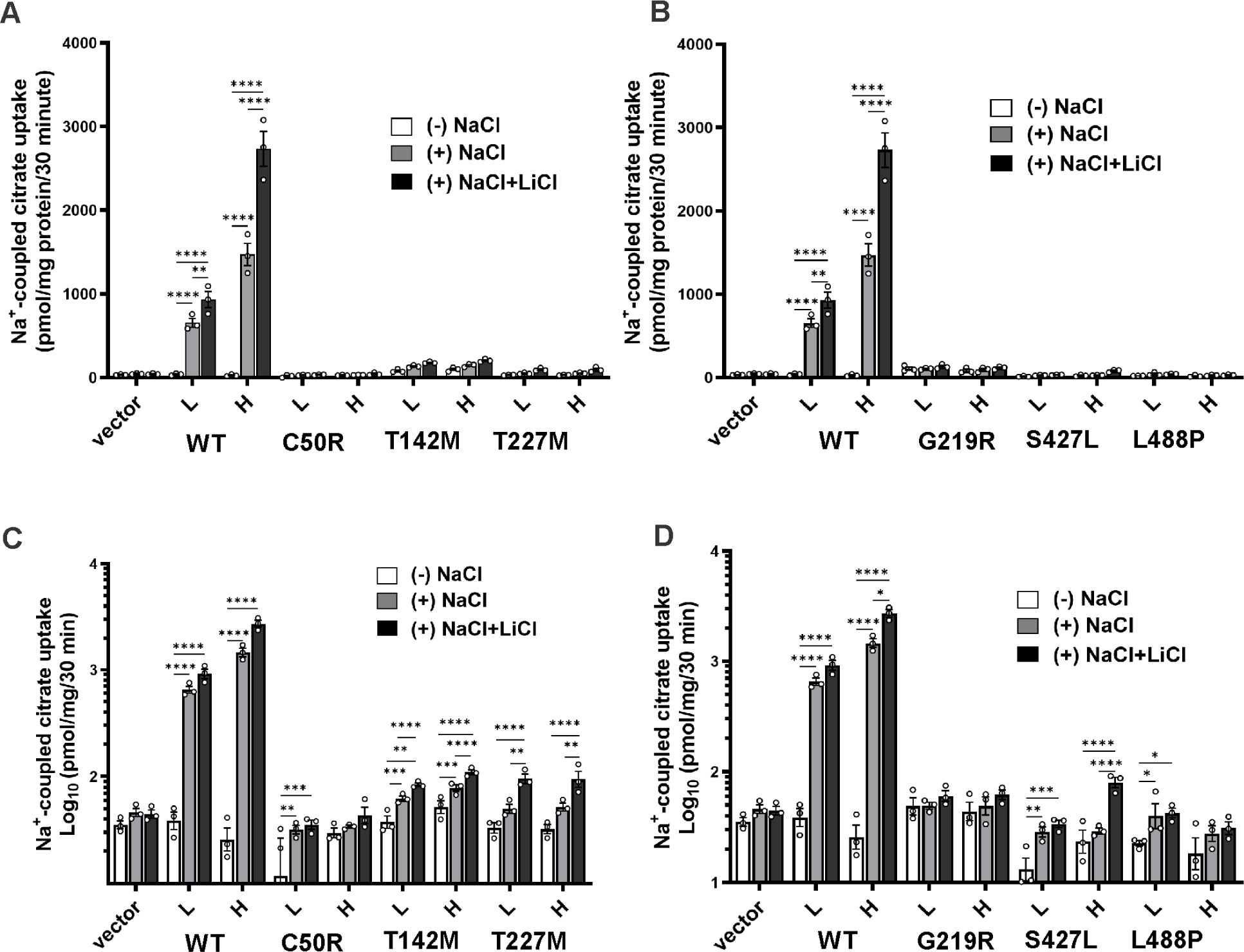
Differential transport activity of WT and mutant NaCTs. Na^+^-coupled uptake of [^14^C]-citrate in HEK293 cells stably expressing (**A**) Class I and (**B**) Class II mutants was measured in buffer devoid of NaCl (white), with NaCl (grey), or with NaCl together with 10 mM LiCl (black) was assayed as described in Methods. (**C**) and (**D**) Log-scale of (**A**) and (**B**) to reveal the very low activity of mutant variants..Each column represents the mean ± SEM, *n*=3-5. \**P* ≤ 0.05, \*\**P* ≤ 0.01, ***P ≤ 0.001 and \*\*\*\**P* ≤ 0.0001.

Since the presence of complex glycosylated proteins infered that the mutant proteins trafficked to the plasma membrane, we independently tested whether the mutants indeed reached the cell-surface and could be biotinylated. Intact cells were treated with the cell-impermeable Sulfo-NHS-SS-biotin that covalently biotinylates surface-exposed primary amine groups of proteins, including NaCT. Biotinylated proteins were subsequently bound to NeutrAvidin resin and separated from non-biotinylated intracellular proteins that do not bind to the resin. **Fig. 2H** and **I** show the results for transiently transfected and clonal cell lines, respectively. In both cases, the upper, complex glycosylated NaCT band was prominent in the bound (B) fractions, indicating that these proteins must have been present at the cell surface at the time of biotinylation. This was clearly visible for WT NaCT and the three mutants C50R, T142M, and T227M mutants, whether transiently transfected (**Fig. 2H**), or expressed at lower or higher levels in clonal cell lines (**Fig. 2I**). The data clearly establish that protein synthesis and trafficking is normal. However these mutant proteins are unable to transport Na^+^/citrate across the plasma membrane (**Fig. 1C**, **F**) likely because either the affinity for substrate is diminished or the transport translocation mechanism is hampered. Based on these data, we categorize C50R, T142M, and T227M as Class I mutations that share similar mechanisms by direclty pertubing the citrate transport functions (Jaramillo-Martinez et al., 2021c).

### Folding mutations do not attain complex glycosylation

First-tier expression analyses of mutants G219R, S427L, and L488P revealed significantly reduced protein expression compared to WT NaCT (**Fig. 4)**. Notably, complex glycosylated proteins (∼90 kDa) were not detectable in cell lysates of transiently transfected cells (**Fig. 4B**, see also quantification in **D**). This was confirmed in 12 clonal cell lines each expressing G219R, S427L and L488P mutants at lower or higher expression levels using the epitope-specific anti-Strep tag antibody (**Fig. 4E-G, Supplemental Fig. S1**). Similar results were obtained when the same samples were developed with anti-SLC13A5-specific antibodies SAB1402084 and HPA044343 (Sigma-Aldrich), or epitope -specific RGS(H)4 (Qiagen). In contrast, antibody PA5-113058 (Invitrogen)(Sun et al., 2023) failed to discriminate between mutants and vector control lanes (**Supplemental Fig. S2**). Again, neither antibodies detected appreciable amounts of the complex glycosylated forms (∼90 kDa) of the mutant proteins.

**Figure 4.**
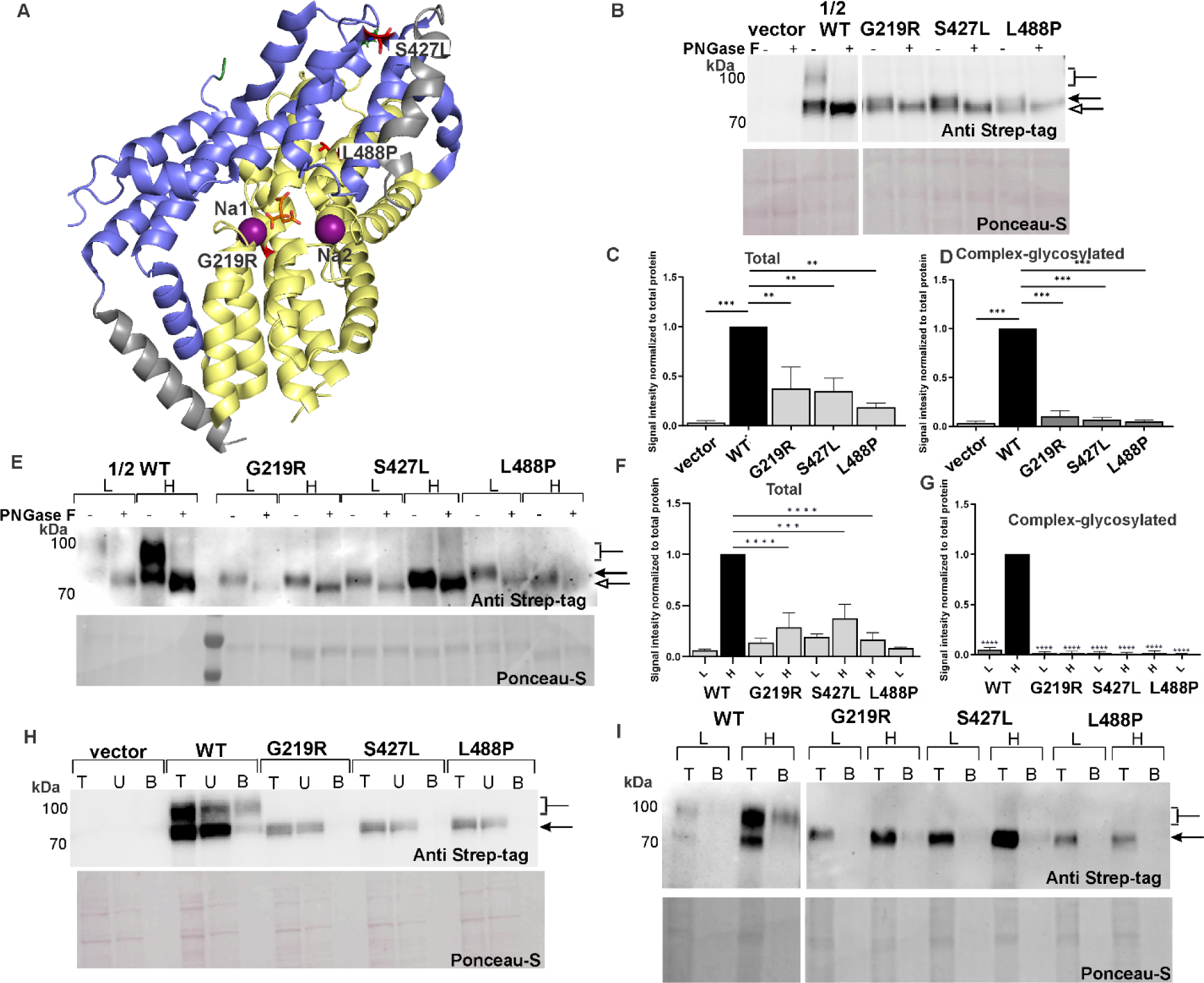
Glycosylation status and expression profiles of Class II mutants. (**A**) Location of G217R, S427L, and L488P in the cryo-EM structure (PDB: 7JSK), colored as in Fig. 1. (**B**) Whole-cell lysates from transient transfections were treated with and without PNGase F as in Fig. 2; note that only 10 µg of WT NaCT was loaded, half as much as mutant cell lysates (20 µg), and resolved by SDS-PAGE and Western blot analysis using the NaCT-specific anti-Strep-tag antibody. Non-glycosylated (empty arrow), core-glycosylated (filled arrow), and complex-glycosylated (brackets) proteins are indicated. (**C**) Quantification of the total Western blot signal of all NaCT bands and (**D**) of the complex-glycosylated band from three independent experiments as decribed in Methods. (**E**) Western blot analysis of whole-cell lysates from clonal cell lines expressing lower (L) and higher (H) level of NaCT variants., and quantification of (**F**) total NaCT signals and the (**G**) complex-glycosylated protein bands. (**H**) Cell-surface biotinylated protein expression in transient transfection and (**I**) clonal cell lines; labeling is as in Fig. 2. Each column represents the mean ± SEM, n= 3. *P ≤ 0.05, **P ≤ 0.01, and ***P ≤ 0.001 The relative total proteins levels were quantified by Image J, n=3.

Closer inspection of the core-glycosylated bands sizes around 75 kDa before and after PNGase F treatment revealed that some lower level glycosylation must have occurred in the mutants G219R, S427L and L488P (**Fig. 4E-G**). If samples were digested with EndoH, an endoglycosydate that selectively cleaves core glycans, the same digestion patterns identical to PNGase F were obatined for the mutant proteins; however, the WT was not accessible to digestion with Endo H (**Fig. 5**). This data suggests that some low-level glycosylations could be attained while the mutants traveled through the ER. Intrestingly, higher overall protein expression in clonal cell line S427L labeled “H” in **Fig. 4E** appeared to correlate with increased lithium-stimulated citrate transport (**Fig. 3B** and **D**).

**Figure 5.**
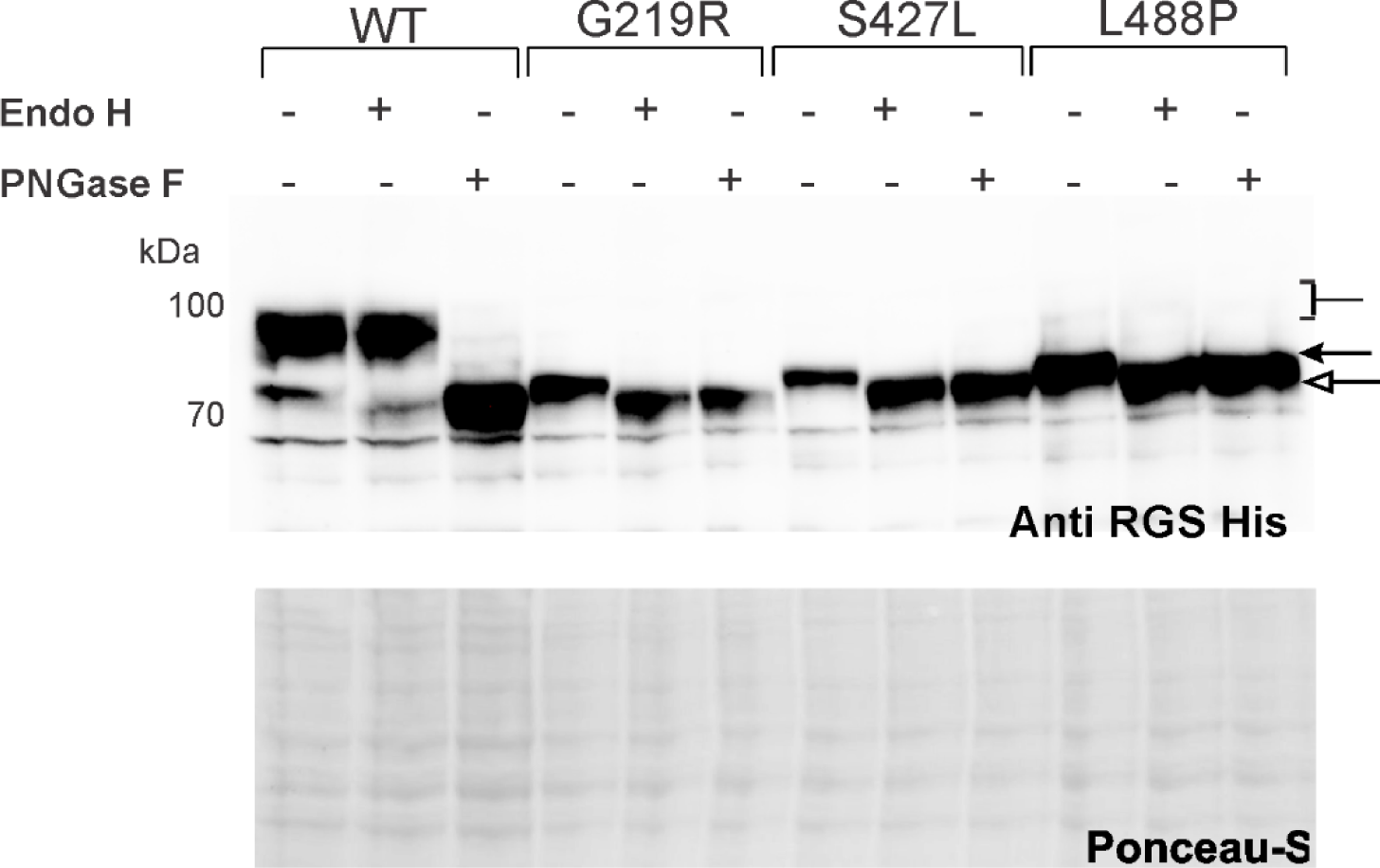
Core and complex glycosylation of WT and Class II folding mutants. Glycosylation patterns of mutants were verified by digestion with Endo H vesrsus PNGase F glucosidases. Endo H is selective for core glycans of the high mannose and some hybrid types of N-linked carbohydrates found in ER proteins. In contrast, PNGase F removes almost all types of N-linked glycosylation of the high mannose, hybrid, bi-, tri- and tetra-antennary types. Note that EndoH did not cleave the complex glycans of WT NaCT, however mutants were readily cleaved, indicating immature ER core-glycosylation.

To confirm that the mutants were unable to traffic to the cell surface, intact cells were treated with the cell-impermeable Sulfo-NHS-SS-biotin, and biotinylated proteins bound to NeutrAvidin resin. Contrary to transport mutants shown in **Fig. 2H** and **I**, mutants G219R, S427L, and L488P completely lacked any detectable signal in the “bound” fraction (B), providing additional evidence that they must not have reached the cell surface. This was true for transiently transfected and stably expressing clones **(Fig. 4H** and **I**). Thus, impaired complex glycosylation could be directly linked to lack of mature protein at the cell surface, which is likely responsible for the lack of Na^+^-coupled citrate transport into cells (**Fig. 3B**). Interestingly, culturing cells at low temperature (25 °C) to slow processing and so extend interaction times with ER chaperones partially rescued their transport activity compared to 37 °C (**Supplemental Fig. S3A**). Together, these criteria classified mutants G219R, S427L and L488P as Class II or protein folding/trafficking mutations.

### Cellular localization of Class II mutants

Based on these results, we aimed to gain further insight in the cellular localizations of WT and mutant proteins by immunocytochemistry. WT NaCT displayed punctuated cell surface staining that co-localized with the plasma membrane marker protein B-catenin **(Fig. 6A, C)**. Clear surface expression was also observed with a GFP-tagged WT NaCT (Jaramillo-Martinez et al., 2022) and is consistent with the robust surface biotinylation observed in **Fig. 2I**. In contrast, the folding mutants G219R, S427L, and L488P predominantly co-localized with Calnexin (**Fig. 6B, D**), a molecular chaperone that assists folding and assembly of nascent proteins in the ER. The Class II folding mutations displayed little if any co-localization with the plasma membrane marker B-catenin. This provided strong evidence that these mutants may interact with ER chaperones; however, they may not escape the ER quality control and therefore cannot travel to the Golgi and plasma membrane, but instead are prematurely degraded. Indeed, no co-localization was found with marker proteins in the Golgi apparatus (GM130),(Briant et al., 2017) lysosome (Lamp1), or endosome (Rab9, **Supplemental Fig. S4**). WT did exhibit some co-localization with Rab9a as is expected for membrane proteins that are internalized by endocytosis and processed back to the cell surface for reuse.

**Figure 6.**
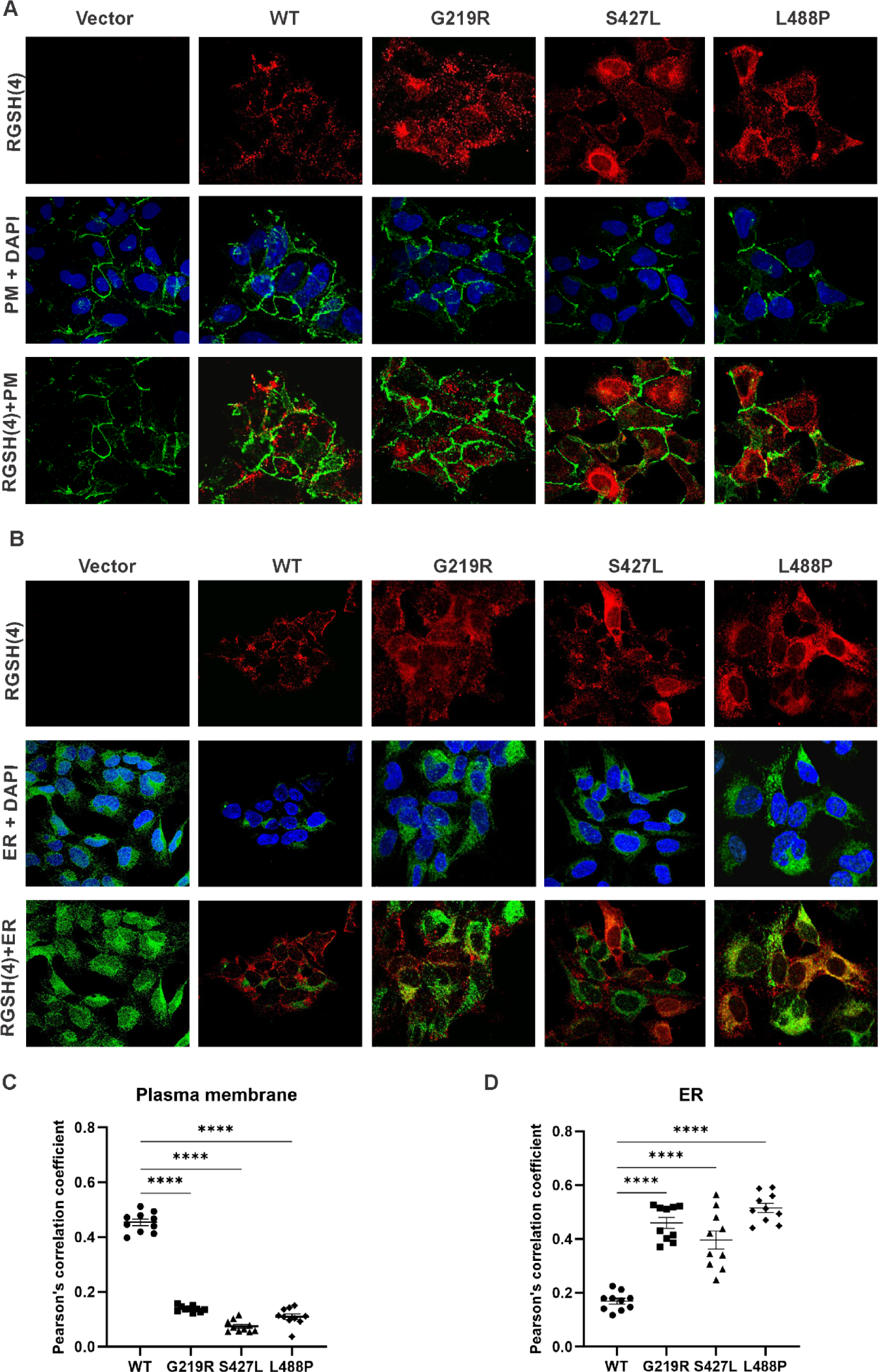
Cellular localization of NaCT folding mutants using multiple cellular markers. Clonal cell lines selected to stably express WT and mutant proteins at lower levels were co-stained with NaCT-specific anti-RGS(H)4 (red) and an antibody recognizing either (**A**) the plasma membrane protein marker B-catenin (green), and (**B)** ER protein marker Calnexin (green). DAPI staining of nuclei was included in blue for reference. (**C**) and (**D**) are Pearson’s correlation coefficient between RGSH(4) and markers. Each point represents an independent image, n=10. *P ≤ 0.05, **P ≤ 0.01, ***P ≤ 0.001 and ****P ≤ 0.0001.

### Half-lives of WT and folding NaCT mutants

To further define the defects of the six common disease mutants, we quantitatively assessed their mRNA levels and studied their mRNA translation and degradation rates. We used real-time quantitative PCR to compare total mRNA levels in cells expressing WT and mutant proteins. The mRNA levels proved similar for most mutants (**Supplemental Fig. S3B**), and somewhat higher for G219R, perhaps to compensate for the phenotype. Similarly, mRNA levels were previously measured by Selch et al. (Selch et al., 2018) with no significant differences detected between mutants G219R, T227M, and L488P. The data exclude mRNA synthesis or degradation as the culprit and reinforce that steps after protein translation into the ER are affected, strongly pointing to protein folding and/or protein stability defects.

To test whether the mutant proteins are less clonal than the wildtype, we studied protein turnover rates. The protein half-life is the time required for half of the protein to be degraded when its regeneration via synthesis is blocked by cycloheximide that halts translation of messenger RNA. Cells were treated with 100 µg/mL cycloheximide (CHX) to halt protein synthesis, and cells were harvested after 0 (control), 4, 8, 12, 16, and 24 h to monitor the decay of the NaCT protein bands. WT retained a strong signal of the complex-glycosylated protein bands for many hours with an estimated half-life of about 23 h (**Fig. 7A, B**). The immature Class II mutant proteins decayed much faster with a half-life of just 3 h observed for S427L, and only 2.8 and 1.6 h for G219R and L488P, respectively (**Fig. 7A, B, bottom panels**). Taken together, not only do these mutants lack complex glycosylated protein, but also is the core-glycosylated immature protein much more labile than WT NaCT, which likely contributes to the detrimental phenotypes.

**Figure 7.**
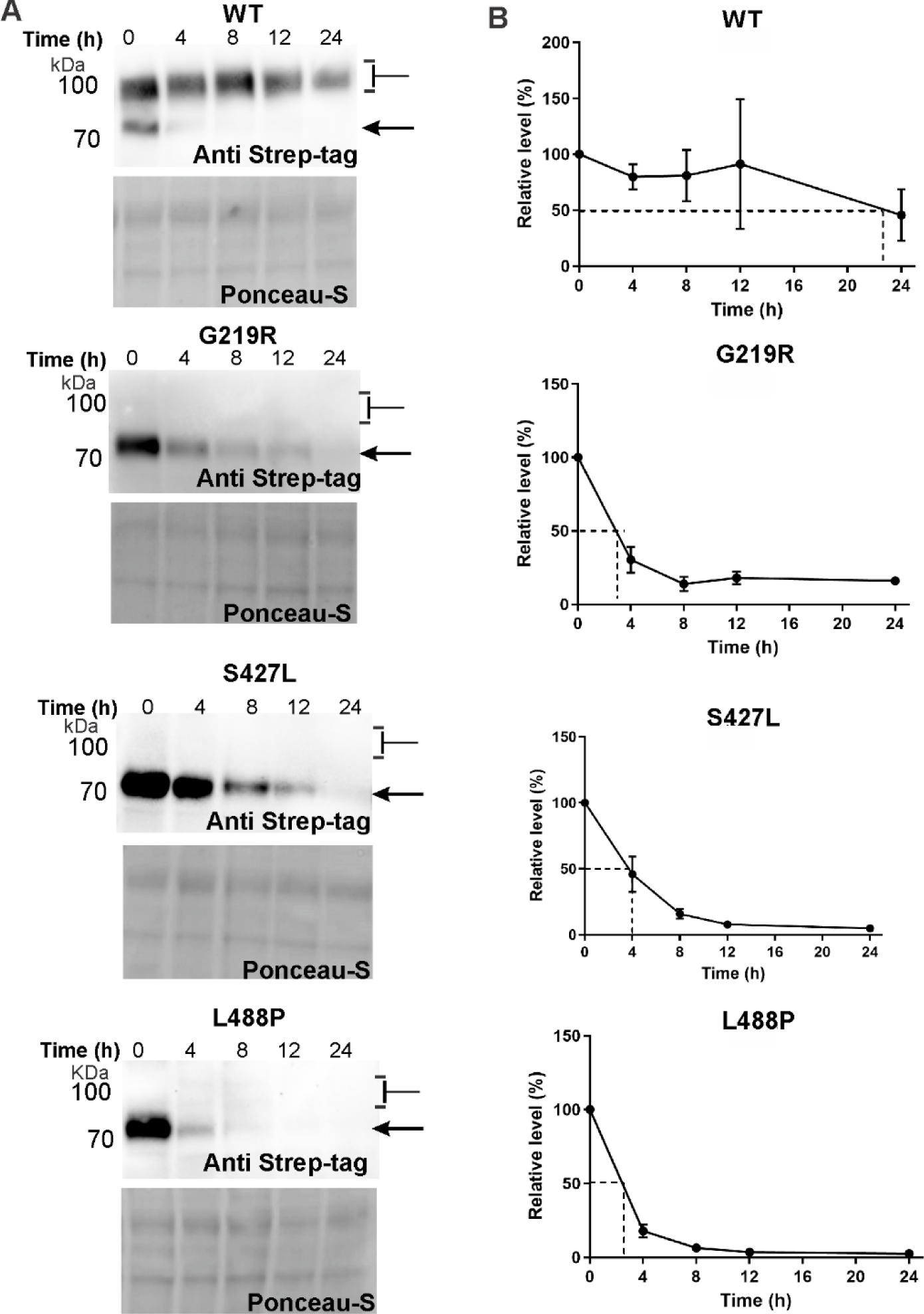
Half-lives of Class II mutants are significantly shortened. (**A)** Clonal cells were treated with cycloheximide, and the disappearance of the NaCT protein bands monitored for 0, 4, 8, 12, and 24 h by Western blotting. Positions of core-glycosylated (arrow) and complex-glycosylated (brackets) NaCT protein bands are indicated. (**B**) Quantification of the total NaCT signals from three independent experiments for the calculation of protein half-lives (see **Table 2**); statistical analyses are described in Methods.

**Table 2.**
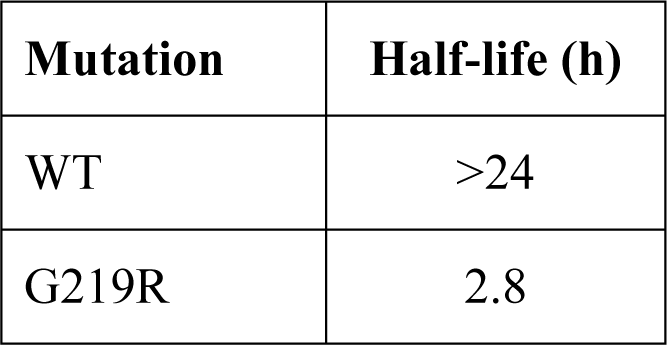
Half-lives of clonal WT and NaCT mutant clones treated with cycloheximide.

### NaCT degradation pathway

To distinguish the degradation pathways of NaCT mutants, clonal cells were treated for 8 h with either proteasomal or lysosomal inhibitors. When MG132, a potent inhibitor of the ubiquitin-proteasome system, was applied, the expression of the immature core-glycosylated band was significantly increased in the mutant proteins, suggesting that G219R, S427L and L488P are degraded in the proteasome (**Fig. 8A** and quantifications in **B**). Further, we verified that total level of ubiquitinated proteins became elevated which is expected when proteasomal degradation is inhibited. **Fig. 8A** (middle panel) showed anticipated response to MG132 treatment for all samples. On the other hand, treatment with the lysosomal inhibitor bafilomycin (Baf) showed no change in the relative expression of the immature core-glycosylated protein bands (**Fig. 8C** and quantifications **in D**). To validate the effectiveness of Baf treatment, we ascertained the elevation of total LC3A/B protein levels. This increase is anticipated when lysosomal degradation is inhibited, as LC3A/B, a ubiquitin-like modifier, participates in the formation of autophagosomal vacuoles that subsequently fuse with lysosomes for degradation. Baf impairs lysosomal function by inhibiting the ability of the vacuolar type H^+^-ATPase (v-ATPase) to transfer protons into the lysosome.(Fedele and Proud, 2020) All of these experiments suggested that proteasomal degradation is the major route for mutants G219R, S427L and L488P.

**Figure 8.**
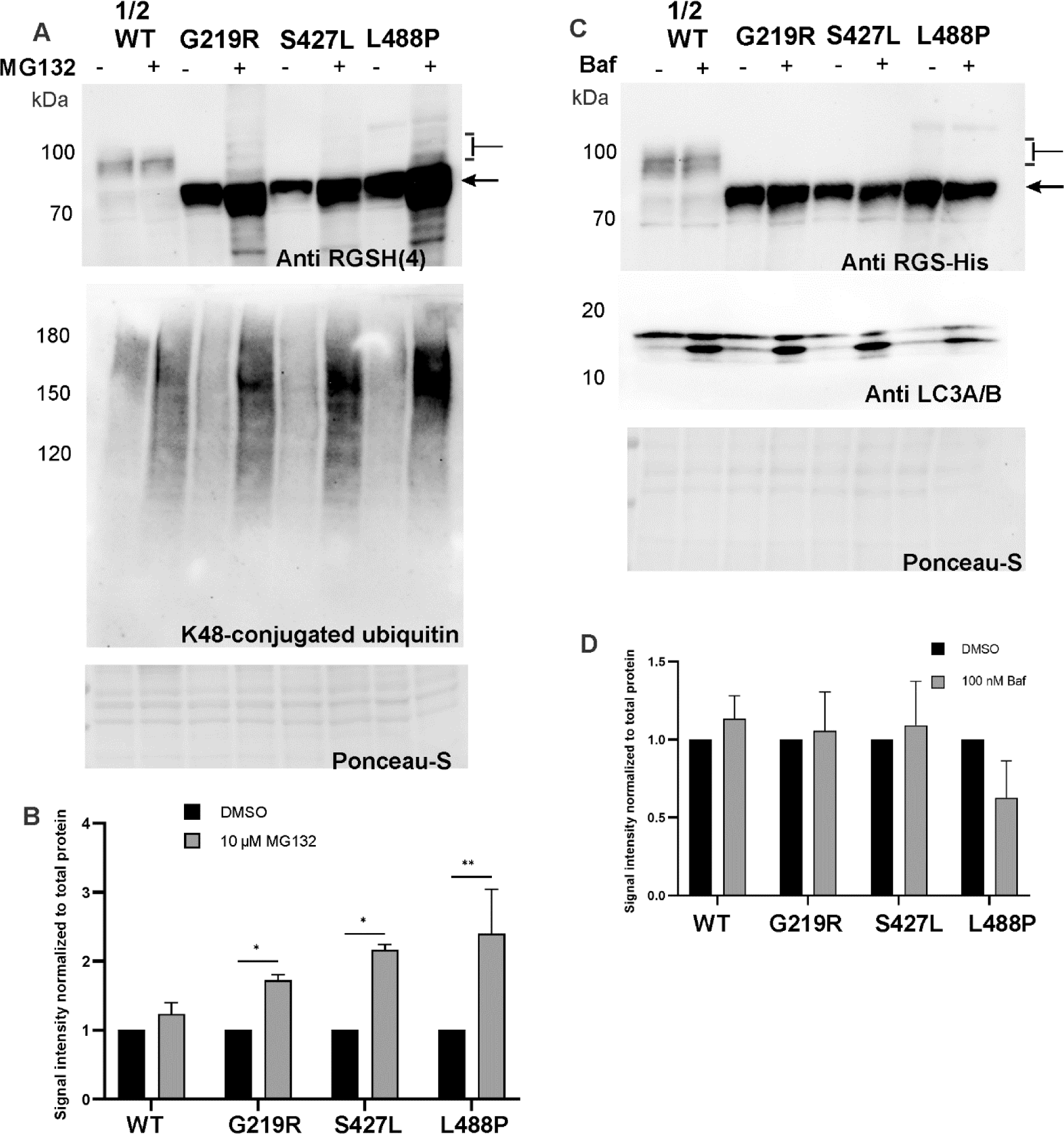
Inhibition of proteasomal and lysosomal degradation pathways for Class II mutants. Clonal cell lines were treated for 8 h with (**A, B**) 10 µM MG132 or (**C, D**) 100 nM bafilomycin A (Baf), and cell lysates analysed by Western blotting. Positions of core-glycosylated (arrow) and complex-glycosylated (brackets) NaCT are indicated. Representative images of 3 experiments are shown, mean ± SEM, n= 3.

## Discussion

### Summary of mutant phenotypes

In this study, we provide biochemical data to elucidate the mechanism of SLC13A5 Epilepsy-causing misssense mutations. All six single amino acid mutations analyzed here severely impaired Na^+^-coupled citrate transport into cells, however, the molecular defects segregated the mutant proteins into two classes. Our findings revealed that Class I mutants C50R, T142M, and T227M traffic normally through the Golgi apparatus, where they attain complete complex glycosylation, as they move to the plasma membrane; they are accessible to surface biotinylation and localize to plasma membrane, similar to WT NaCT (**Fig. 2**). Our findings also revealed that **Class II mutants G217R, S427L,** and **L488P** only attain immature core glycosylation, and cannot be biotinylated when intact cells are exposed to a membrane impermeable biotinylation reagent, strongly indicating that they are retained intracellularly and do not reach the plasma membrane (**Fig. 3**). The sizes of the immature poteins were verified with two different NaCT-specific epitope-tag antibodies and confirmed with two anti-SLC13A5 antibodies. Their distinct core-glycosylation patterns and succeptability to cleavage by both PNGaseF and EndoH suggest that the proteins are retained in the ER (**Fig. 5**). This was confirmed by immunohistochemistry demonstrating distinct co-localization with ER maker Calnexin (**Fig. 6B**). Together, the data strongly suggest minimal escape of these mutant proteins from the ER quality control. This was supported by low temperature rescue which partially restored citrate transport in the order L488P > S427L > G217R. Noticably, mRNA levels were similar in mutants and WT NaCT. The significantly shorter turnover rates of the core-glycosylated mutant proteins (**Fig. 7**) further indicated protein folding, posttranslational modification, or protein stability defects, and premature degradation by the ER quality control machinery that feeds the proteasomal degradation pathway.

### Phenotypes confirmed in other studies

The data are, in principle, in line with earlier reports by Hardies et al. (Hardies et al., 2015) on the T142M, T227M, G219R and S427L mutations and by Selch et al. (Selch et al., 2018) on the T227M, G219R and L488P mutations with some caveats. Hardies et al. reported expression profiles for WT and transport mutations T142M, T227M in HEK293 cells with primary, secondary and tertiary protein bands between 49 and 40 kDa that are much smaller than the calculated molecular weight of the V5-tagged protein of 64 kDa. Moreso, in Western blots of folding mutations S427L and G219, only the lower secondary band around 40 kDa was detectable which may reflect their greater succeptibility to proteolytic degradation. The heavily degraded proteins and absence of full-length, immature or mature NaCT proteins in that study are of course a concern. The authors did employ an ER marker KDEL for immunocytochemistry to colocalize the two folding mutations S427L and G219 to the ER. On the other hand, the study by Selch et al. showed expression of all mutants and WT with the same band pattern around 65 kDa in total membrane fractions, suggesting full-length proteins (calculated size of untagged NaCT is 63 kDa) but with a distinction that G219R and L488P were not detectable in the isolated plasma membrane fraction. The absence of the G219R and L488P mutant proteins in plasma membranes was corroborated by immunofluorescence analyses albeit no marker proteins were used to affirm their localization to the ER. Thus in principle, both studies agree with our classification of the T142M and T227M as transport mutations (Class I), and G219R, S427L and L488P as folding mutations (Class II). Our current study is the first to report on C50R as a transport mutation (Class I).

In contrast, the report by Klotz et al. (Klotz et al., 2016) on the T227M, G219R and L488P mutations is at odds to our data as well as Hardies’ and Selch’s. Klotz et al. reported for the G219R mutation in transiently transfected monkey kidney COS-7 cells a full-length protein of about 75 kDa that was found at the cell surface in biotinylation experiments, and therefore was characterized as transport mutation. This group found a small amount of citrate transport activity in G219R transfected COS-7 cells, but not HEK293 cells, perhaps pointing to cell-type specific variability of background activity. Other discreancies are the lack of detectable NaCT protein in Western blots of the T227M and L488P mutations that therefore were both characterized as folding mutation. It is not clear why T227M was not detected in that study. In our experience, folding mutations including L488P were expressed at much lower levels than WT NaCT in the order WT>>S427L>G219R>L488P, the latter was only detected after prolonged exposure and gave a weak signal compared to WT even though only half or a third of WT protein was loaded (**Fig. 4, Supplemental Fig. S1E-G**).

Notably, none of the above studies probed the stimulatory effect of lithium on the sodium-coupled citrate transport that is specific for the NaCT transporter in humans and primates (Gopal et al., 2015; Inoue et al., 2003). Here, we demonstrate that lithium, at concentrations employed for treatment of bipolar disorders, enhanced the low-level activity of several mutant proteins C50R, T142M and L488P mutants suggesting that Li may be incorporated in treatment strategies for these mutants.

### Glycosylation status and ER associated protein quality control

Furthermore, none of the prior studies used endoglycosidases to probe the glycosylation patterns of the proteins and distinguish immature proteins retained in the ER (cleaved by Endo H) from complex glycosylated proteins (cleaved by PNGase F) that have maturated while trafficking through the Golgy compartment (**Fig. 5**). Such glycosylation analyses are routine for phenotyping of CFTR folding mutations leading to cyctic fibrosis (Grove et al., 2011a; Grove et al., 2011b; Hildebrandt et al., 2014; Meacham et al., 1999; Okiyoneda et al., 2013; Patrick et al., 2011; Rubenstein et al., 1997; Rubenstein and Zeitlin, 1998; Taylor-Cousar et al., 2019; Yang et al., 2018) and, as we show here, are a simple and an effective tool for future probing of additional SLC13A5 Epilepsy causing mutations. Moreover, most previous publications on SLC13A5 Epilepsy-causing mutations relied on transient transfections (Klotz et al., 2016; Selch et al., 2018) which give inherently non-homogenious populations of NaCT expressing cells. Here, we generated clonal cell lines stably expressing lower and higher levels of NaCT mutants to ascertain that results are independent of the amounts of protein produced in cells. Interestingly, the WT and Class I NaCT proteins displayed abundant complex-glycosylated protein and little of the core-glycosylated band sizes. In contrast, transiently transfected cells exhibited similar amounts of the complex- and core-glycosylated proteins, perhaps because of a relatively short expression window of just 48 h, or perhaps due to overexpression and “overloading” of the expression machinery in a few positive cells. Importantly, in both cases the core-glycosylated bands when treated with PNGase F collapsed into an even lower size band suggesting that some short-chain N-linked glycans were added to Asp562 in NaCT as expected when trafficking through the ER (Schoberer et al., 2018; Wang et al., 2015). The clonal NaCT mutants’ cell lines may offer more consistent and reproducible results for future large-scale drug-screening efforts.

To further analyse the defects of Classs II mutants, we studied their protein turnover rates, mRNA levels, and protein degradation pathways. G219R, S427L and L488P displayed significantly shorter half-lifes compared to WT (**Fig. 7**), indicating that degradation of the immature protein in the ER occured at a much accelerated pace. At the same time total mRNA levels were mostly similar among WT and mutants indicating similar mRNA translation and degradation rates, with G219R showing a slight increase, possibly compensating for the phenotype (**Supplemental Fig. S3B**). We conclude that the pathology of the mutations is unlikely related to mRNA. Misfolded proteins that are retained in the ER are subsequently degraded either by the proteasome through the ubiquitin-dependent endoplasmic reticulum-associated degradation pathway (ERAD), or via the lysosomal degradation pathway (Katayama et al., 2015; Thévenod and Friedmann, 1999; van Kerkhof et al., 2001; Wei et al., 2005; Williams et al., 2013). The proteasome inhibitor MG-132 increased the accumulation of immature mutant proteins in the ER (**Fig. 8A, B**) but did not increase trafficking to the Golgi since it did not restore the folding imbalance suggesting that proteasomal degradation is the major route of degradation for G219R, S427L, and L488P. Taken together, our results lay out an experimental roadmap to phenotype other SLC13A5 Epilepsy causing mutations, and set the stage for small molecule screening efforts to begin.

Finally, we present here, for the first time, a fluorescence tranport assay that relies on a high-performance genetically encoded citrate biosensor (Citron) for the detection of intracellular citrate accumulation. The assay is safer than current [^14^C]-citrate uptake assays, and displayed a robust, rapid increase of fluorescence within minutes in WT NaCT, yet was sensitive enough to detect residual low-level activity in one of the folding mutations L488P. The assay can be performed in multi-well formats and may open up the door for future discovery screening of small molecules therapeutics.

### Significance

The lack of transport function of NaCT has a significant impact in neuronal cells that require citrate as key metabolic intermediate to boost their energy needs, and to serve as a precursor for the synthesis of cholesterol and fatty acids (Gopal et al., 2007) as well as the neurotransmitters acetylcholine, GABA, and glutamate (Yodoya et al., 2006). Any imbalance in the three neurotransmitters may be the underlying cause of epileptic episodes observed in SLC13A5 Epilepsy patients in addition to a general lack of sufficient energy from citrate import leading to frequent epileptic encephalopathies and intellectual disabilities observed in patients (Schossig et al., 2017). The same can be speculated in terms of the loss of transport function of NaCT in astrocytes. Cytoplasmic citrate is likely to play an important role in the glutamine-glutamate cycle where astrocyte-associated glutamine synthesis/release is a salient component of the cycle. Citrate can be converted to glutamate via the actions of aconitase-1, isocitrate dehydrogenase IDH1, and glutamaite dehydrogenase for subsequent conversion to glutamine. The glutamine-glutamate cycle that operates between astrocytes and glutamatergic neurons is critical for cognitive development. As such, the intact function of NaCT in neurons and astrocytes is obligatory for normal development of brain functions.

### Structural implications of mutations in the context of the local environment

In the 3D-structure (PDB: 7JSK (Sauer et al., 2021a)), residues T142 and T227 (**Fig. 2A**, green sticks) participate in hydrogen bonding to citrate (orange sticks). Substitution of the threonine sidechain with a methionine sidechain will disrupt these binding interactions and significantly reduce binding affinities for the substrates (Jaramillo-Martinez et al., 2021a; Jaramillo-Martinez et al., 2021c; Mancusso et al., 2012; Sauer et al., 2021a). On the other hand, residue C50 is located at the interface between the scaffold (blue) and transport domains (yellow). Substitution with a large, positively charged arginine will significantly decrease hydrophobicity (**Supplemental Fig. S5A**) and may, through steric interactions, hinder movement of the two domains and thus transport. In any of these Class I mutants, identifying “potentiator”-like molecules that somehow increase transport may be a future treatment option. Such potentiators have been highly successful in the treatment of gating mutations of CFTR (e.g., G551D); they act by non-specifically opening the channel to enable ion fluxes. However, potentiators have not been described so far for any disease-causing mutations of transporters that utilize an elevator-type mechanism, such as NaCT (Jaramillo-Martinez et al., 2021a; Jaramillo-Martinez et al., 2021b; Jaramillo-Martinez et al., 2021c; Mulligan et al., 2016; Sauer et al., 2021a; Sauer et al., 2021b).

Class II folding mutations are dispersed throughout the NaCT structure, with G219R residing about >5 Å away from the closest sodium ion (Na1, magenta ball in **Fig. 4A**) too far away to directly participate in Na-citrate binding. It is located at the end of helix TM 5a that together with inverted helix TM 10a is critical for forming the substrate-binding pocket. Substitution of glycine with large, positively charged arginine will significantly decrease hydrophobicity and may, through steric interactions, distort the binding site. This is supported by predicted stability changes (ΔΔG) computed by several web servers (**Supplemental Fig. S5**) and is in agreement with the destabilizing effect on protein previously predicted by Hardies et al. (Hardies et al., 2015). Substitution of L488P in the C-terminal transport domain (yellow) may cause location distortion of α-helical structure by helix breaker proline; this is likely responsible for folding defects. L488P decreased hydrophobicity of surrounding residues, and was largely destabilizing (**Supplemental Fig. S5B**). Substituting polar serine S427 with hydrophobic leucine may increase hydrophobicity of surrounding residues and affect α-helix packing, thereby preventing the protein to propely fold. Predicted stability changes at this position gave contradictory results and were unreliable, and underline the notion that phenotypes need to be determined experimentally.

### Rescue strategies for Class II folding mutations

The Class II folding mutants may be capable of actively transporting citrate into cells, if they were able to reach the plasma membrane. In analogy to other plasma membrane transporters, newly synthesized plasma membrane proteins, as they emerge from the ribosome and are inserted into the ER through the SEC61 translocon, attain N-glycosylation and undergo co-translational domain folding and insertion into the membrane facilitated by the chaperone calnexin. For NaCT, proper assembly of the multiple scaffold-scaffold and scaffold-transporter domain interfaces likely will be critical in the assembly process for the transporter to pass the ER quality control checkpoints, and to traverse the Golgi complex where the native NaCT will undergo complex-glycosylation and reach the plasma membrane by vesicular trafficking. Therefore approaches to correct the protein folding defect and facilitate escape from the ER maybe feasable as future treatment options. Small molecule “correctors” have been highly successful for treatment of the folding defect of the prominent CFTR-F508del mutant that causes severe cystic fibrosis (Grove et al., 2011a; Grove et al., 2011b; Hildebrandt et al., 2014; Meacham et al., 1999; Okiyoneda et al., 2013; Yang et al., 2018).

### Conclusions

Our model in **Fig. 9** proposes that Class I mutations traffick normally to the cell surface but cannot transport citrate inside the cells to satisfy the demand for citrate in the cytoplasm. Class II mutations, upon co-translational folding and insertion into the ER membrane, cause local misfolding of the protein that is recognized by the ER quality control machinery and the protein is targeted for degradation. This opens an opportunity for domain-specific targeting of small-molecule therapeutics to correct the folding defects, thereby promoting trafficking to the cell surface to perform the transport function. Meanwhile, gene therapy may also be explored to correct the SLC13A5 disease-causing mutations. Notwhithstanding, these approaches however rely on thorough understanding of the mechanisms caused by the underlying genetic defect of individual mutations or of classes of mutations with a similar phenotype. The data provided here may provide a roadmap for analyzing additional SLC13A5 Epilepsy mutations and provide a tool set to guide further developments towards strategies to rescue transport functional defect of Class I mutations and to correct protein folding/trafficking defects of class II mutations.

**Figure 9.**
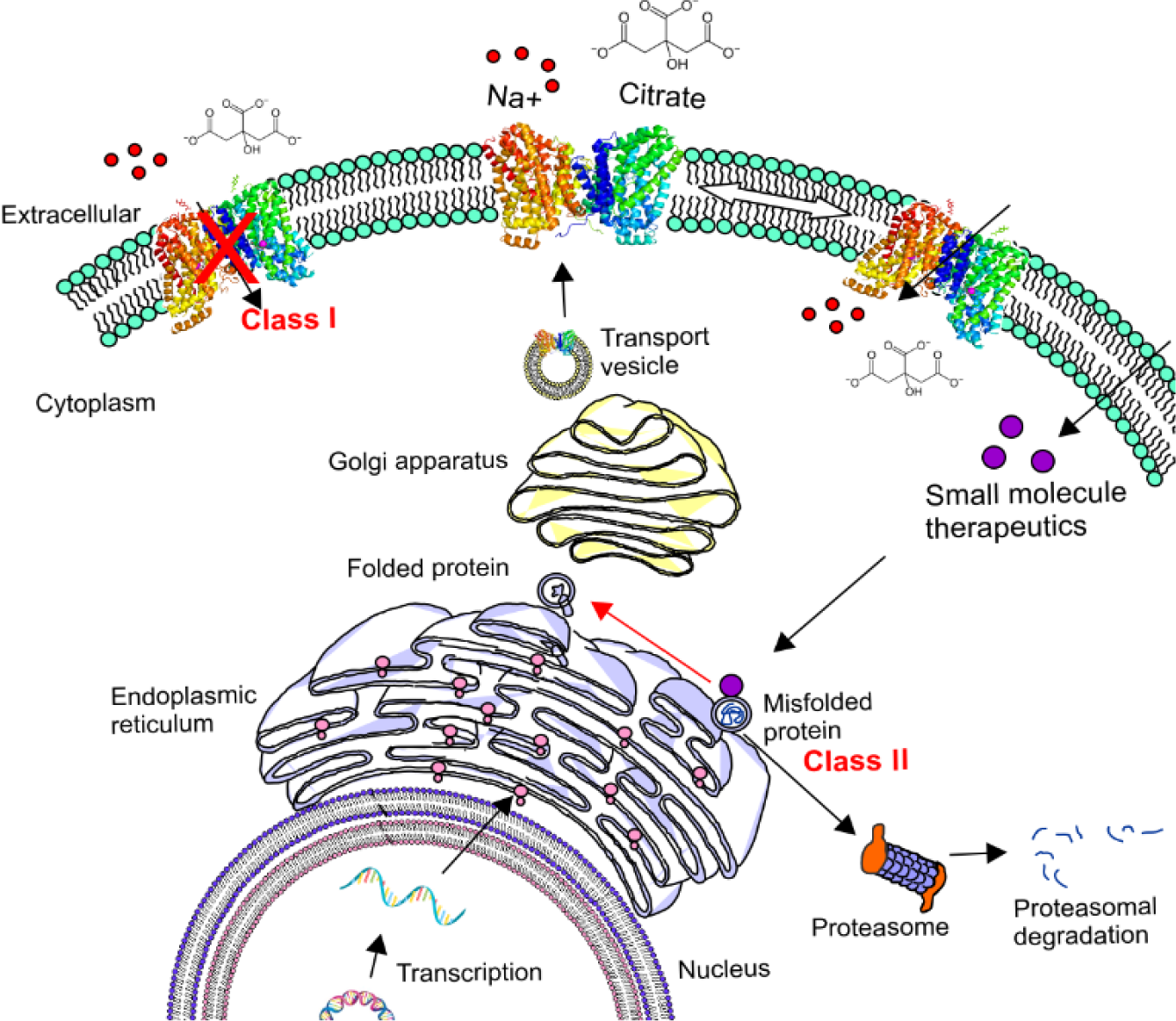
Phenotypes of disease-causing NaCT missense mutations. Based on our experimental analyses, disease-causing missense mutations either (i) interfere with the citrate transport function (Class I) by blocking critical steps in the substrate translocation or (ii) cause defects in protein folding and trafficking (Class II) and lead to premature degradation by the ER quality control machinery. Potential treatment strategies for Class II mutations may include small molecule therapeutics that traverse the cell membrane and reach the ER where they bind to the mutant proteins and ‘correct’ the folding defect and rescue the immature protein, thus promoting it to traffic to the plasma membrane. Potential treatment strategies for Class I mutations may include a different kind of small molecules that may bind to the mutant transporter allosterically and ‘potentiate’ symport of Na^+^ and citrate into the cytoplasma.

## Materials and methods

### Materials

Sulfo-NHS-SS-Biotin was purchased from ThermoFisher Scientific. [^14^C]-Citrate (specific radioactivity, 113 mCi/mmol) was purchased from Moravek Biochemicals. Cycloheximide, carbobenzoxy-Leu-Leu-leucinal (MG132), and Bafilomycin A1 (Baf) were purchased from Millipore-Sigma (St. Louis, MO, U.S.A.). All other reagents were from Millipore-Sigma (St. Louis, MO, U.S.A.) or Thermo Fisher Scientific (Waltham, MA, U.S.A.).

### Plasmids with DNA mutations

To facilitate detection of NaCT, we engineered N-terminal epitope tags RGSH_10_, TwinStrep, and SUMOstar (named HSS*-NaCT, NCBI gene accession number MZ367587).(Jaramillo-Martinez et al., 2022) The calculated molecular weight of NaCT including the tags ∼81.23 kDa. Mutants C50R, T142M, G219R, T227M, S427L, and L488P were introduced by site-directed PCR mutatgenesis using Phusion II hotstart DNA polymerase (Thermo Scientific). Primers with diagnostic sites are listed in Supplemental **Table 1**. Open reading frames of all mutant genes were confirmed by DNA sequencing.

### Cell lines and transient transfection

Human embryonic kidney HEK293-FT cells (RRID: CVCL_6911, Catalog number: R70007) carrying the T antigen of SV40 on a neomycin resistance were purchased from ThermoFisher Scientific. HEK293-FT cells were cultured in high-glucose DMEM medium supplemented with 10% FBS and 1% penicillin/streptomycin at 37 °C under 5% CO2. HEK293-FT cells were grown in 24 or 12-well collagen-coated tissue culture plates to 80% confluency, and transiently transfected with pcDNA 3.1(-) carrying either the WT HSS*-NaCT gene or mutations C50R, T142M, G219R, T227M, S427L, and L488P, or the “empty” pcDNA 3.1(-) vector with Lipofectamine-3000 following established protocols (Jaramillo-Martinez et al., 2022). The cells were allowed to recover for 48 h before assays.

### Generation of clonal cell lines

To establish cell lines stably expressing NaCT variants, HEK293 (lacking the T antigen of SV40, purchased from ATCC, Catalog CRL-1573) were transiently transfected with WT or the mutant genes C50R, T142M, G219R, T227M, S427L, and L488P as described above, and cultured in EMEM medium supplemented with 10% FBS and 1% penicillin/streptomycin. 48 h post transfection, cells were diluted and selected in media containing 500 µg/ml Geneticin®, taking advantage of the neomycin resistance gene of the pcDNA3.1 vector. Monoclonal colonies (about 10 per mutant) stably expressing the exogenous NaCT were isolated, expanded, and cryopreserved at different passages of the selection process (Chaudhary et al., 2012; Chaudhary et al., 2016). All cell lines were used between passages 3 and 13 for the experiments.

### Western blot analysis

For Western blot analyses, cells were lysed with RIPA buffer (25 mM Tris/HCl pH 7.4, 150 mM NaCl, 1% Triton X-100, 1% sodium deoxycholate, 0.1% SDS) supplemented with protease inhibitors leupeptin, pepstatin A, chymostatin and PMSF for 30 min on ice, and total protein concentrations of cleared cell lysates were measured with the Pierce^TM^ BCA Protein Assay Kit (ThermoFisher Scientific) using BSA as a standard. 20 µg of total proteins were dissolved in Laemmli buffer contaning 10% SDS, incubated at room tempeture for 10 min, resolved on 10% or 4 – 15% acrylamide gradient gels together with PageRuler prestained protein ladder (ThermoFisher), and transferred to nitrocellulose membranes. Heating the samples above 60 °C led to membrane protein aggregation. For the detection of NaCT, anti-RGS(H)_4_, anti-Strep-tag II, and anti-NaCT antibodies were used as indicated with the SuperSignal West Pico PLUS Chemiluminescent substrate (ThermoFisher Scientific). For detection of K48 and LC3, anti-Ubiquitin and anti-LC3B antibodies were used; dilutions and catalog numbers are listed in **Supplemental Table 2.** Western blot signals were recorded with ChemiDoc MP Imaging System (Bio-rad Laboratories Inc.) and protein band intensities quantified using ImageJ software (Rasband, 2011). Total proteins loaded per lane were visualized with Ponceau S staining and quantified using Image J software.

### Uptake measurement

[^14^C]-Citrate uptake was measured in the presence of 140 mM NaCl for 30 min at 37 °C. In the absenc of NaCl, 140 mM N-methyl-D-glucamine chloride (NMDG-Cl) was added to maintain equimolarity of chloride ions. LiCl (10 mM) was added to Na^+^-containing samples to stimulate NaCT-specific citrate transport as previously described (Jaramillo-Martinez et al., 2022). Uptake values were normalized to the protein content of lysed cells.

### Citron transport assay

HEK293FT cell were transiently co-transfected with the fluorescent citrate sensor Citron (CMV-Citron1, AddGene #134303 (Zhao et al., 2020)) and either pcDNA3.1 carrying the WT or mutant genes T227M, S427L, and L488P as described above, and grown for 36-48 h to about 90% confluency. Cells were washed twice with transport buffer containing 140 mM NaCl, and fluorescence (excitation 488 nm, emission 517 nm) was recorded every 2 min in a fluorescence plate reader (Tecan Infinite® 200 PRO). At indicated times (see arrows), 10 mM citrate was added, followed by 10 mM Li^+^. To reveal the total possible fluorescence, which depends on Citron expression levels, 0.5% of the mild detergent digitonin was added to permeabilize the cells and allow citrate to enter freely.

### Cell-surface biotinylation

For cell-surface biotinylation, cells were typically grown in 12 well tissue culture plates until they reached 80% confluency. Cells were washed with PBS and exposed to 0.5 mg/mL membrane-impermeable EZ-Link Sulfo-NHS-SS-biotinylation reagent (ThermoFisher Scientific) for 20 min at 4 °C. The reaction was quenched by three washes with 200 mM glycine in PBS. Cells were lysed with RIPA buffer as described, and the lysate supernatant incubated with NeutrAvidin overnight at 4 °C (50 µl packed resin per well). The resin was washed with 20 column volumes of RIPA buffer to remove unbound proteins. Biotinylated proteins were eluted from the resin with one column volume of Laemmli buffer contaning 10% SDS. Samples (30 µl) were resolved by SDS-PAGE for Western blot analyses to semi-quantitatively access the biotinylated fraction of NaCT that bound and was eluted from the resin (labeled “B”) compared to the total NaCT content of cell lysates (“T”) versus unbound NaCT in the wash (“U”) (Date et al., 2016; Huang, 2012; Patrick et al., 2011).

### Immunofluorescence assay and confocal microscopy

Cells were seeded at a 50% confluency onto collagen-treated coverslips and treated as described in (Jaramillo-Martinez et al., 2022). Briefly, cells were washed with PBS, fixed with 4% paraformaldehyde for 15 min, washed with PBS, permeabilized with 0.1% Triton X-100 for 5 min and blocked with 1% BSA for 1 h. The coverslips were co-incubated with mouse anti-RGS(H)_4_ and one other primary antibody (derived from rabbit) recognizing a marker protein typically localized either in the plasma membrane (PM), endoplasmic reticulum (ER), Golgi apparatus (Golgi), endosomes, and lysosomes for 1 h with appropriate dilutions (**Supplemental Table 2)**. The next day, coverslips were washed with PBS and incubated with goat anti-mouse Alexa Flour 647 and goat anti-rabbit Alexa Fluor 488 secondary antibodies and mounted using ProLong Diamond Antifade Mounting with DAPI. The images were visualized at 23°C using a scanning confocal microscope Nikon T1-E with a 60x objective and analyzed using NIS software (Nikon Instruments Inc.). The images represent a maximum projection intensity derived from a Z-stacks.

### Real-time quantitative PCR

RNA was purified from transiently transfected HEK293-FT cells by NucleoSpin RNA (Takara) and the concentrations determined by Nanodrop One (ThermoFisher Scientific). The cDNA was prepared by High-Capacity cDNA Reverse Transcriptase kit (Applied Biosystems). Real-time-qPCR primers for SLC13A5 and control GAPDH are listed in **Supplemental Table 2**. The details of the real-time-qPCR reaction using Power SYBR Green PCR Master Mix (Applied Biosystems) in a Quant Studio 12K Flex Real-Time PCR machine 285880228 (Applied Biosystems) have been described previously.(Karamysheva et al., 2018) The mRNA levels of the SLC13A5 gene variants were calculated using the ΔΔCt method (Schmittgen and Livak, 2008). First, the SLC gene Ct value was normalized to the GAPDH (control gene) (ΔCt) and then to the WT SLC13A5 gene level (ΔΔCt).

### Pharmacological treatment

To assess the half-life of wildtype NaCT and mutant proteins, stably expressing cells were incubated with 100 ug/mL cycloheximide for 0, 4, 8, 12, and 24 h. To distinguish between proteasomal and lysosomal pathways in clonal cell lines, the cells were incuabated with inhibitors carbobenzoxy-Leu-Leu-leucinal (MG132) or Bafilomycin A1 (Baf) for 8 h at 10 µM and 100 nM respectively.

### Statistical Analysis

In Western blot analysis, the ECL signal intensity of the NaCT protein bands was quantified using ImageJ (Rasband, 2011). To account for variations, each band was normalized with the Ponceau S stain signal, which represents the total protein loaded per lane. Means ± SEM of three to four independent experiments were calculated and graphed with the GraphPad Prism 9 Software (San Diego, California, USA). To assess statistical differences between control and experimental groups, two-tailed unpaired Student’s t-tests were used followed by Dunnett’s test of multiple comparisons. Significance was set at *P* < 0.05. [^14^C]-Citrate uptake measurements into whole cells were conducted in triplicate wells, and normalized to total protein of cell lysates found in each well. To assess statistical differences between control and experimental groups, one-way analysis of variance followed by multiple comparisons Tukey’s test was used. Significance was set at *P* < 0.05. WT, mutant variants and controls were assayed in parallel in triplicate wells. For citron assay, statistical analyses of at least three independent experiments were performed with the GraphPad Prism 9 Software. The half-lives of WT and mutant NaCTs were calculated using the One Phase decay equation: *Y*0 ∗ exp(-*K* ∗ *X*) Y, where *Y0* represents the value of Y at zero time, *K* is the rate constant (reciprocal of the time units on the X-axis), and X-axis units correspond to half-life (computed as ln(2) /*K*). The Pearson’s correlation coefficient was calculated for 10 independent images. The statistical differences were analyzed by One-Way Anova followed by Dunnett’s test of multiple comparisons.

## Supplementary material

Table S1 details the primers for site-direct PCR mutagenesis and for real-time quantitative PCR. Table S2 lists the antibodies used in immunofluorescence assays, along with specific dilutions, supplier information, and catalog numbers. The following supplemental figures are included. Fig. S1: NaCT expression levels in clones selected to stably express NaCT WT, C50R, T142M, T227M, G219R, S427L and L488P. Fig. S2: Comparison of NaCT specific antibodies. Fig. S3: Low temperature partially rescues transport activity of mutants with folding defects and mRNA levels are unchanged in NaCT mutants. Fig. S4: Cellular localization of NaCT folding mutants. Fig. S5: Theoretical calculations of the impacts of disease-causing mutations on hydrophobicity and protein stability.

## Data availability

The data will be made accessible by the corresponding authors upon the submission of a reasonable request.

## Acknowledgements

We thank Qinghai Zhang for critical reading of the manuscript and insightful comments. We are grateful for the support of an Early Career Investigator Research Grant from the TESS Research Foundation to VJM. The work was supported, in part, by National Institutes of Health grant GM141216 to ILU. Author contributions: V. Jaramillo-Martinez: Conceptualization, Data curation, Formal analysis, Funding acquisition, Investigation, Methodology, Validation, Visualization, Writing—original draft, Writing—review & editing; S.R. Sennoune: Investigation, Writing—review & editing; D.A. Agard: Methodology, Supervision, Writing—review & editing; E.B. Tikhonova: Investigation, Writing—review & editing; A.L. Karamyshev: Writing—review & editing. V. Ganapathy: Writing—review & editing. I.L. Urbatsch: Conceptualization, Funding acquisition, Project administration, Resources, Supervision, Validation, Writing—review & editing.

## Authors Noters

Disclosures: The authors declare no competing interests exist.

