## Supplementary Materials for "Molecular phenotypes segregate missense mutations in SLC13A5 Epilepsy"

### Supplementary material

**Supplemental Table 1 Primers for site-direct PCR mutagenesis and for real-time quantitative PCR.** The base pairs and codons changed are shown in bold and underlyned, respectively. All primers were ordered from Sigma Aldrich.

| **Mutation** | **Primers** | **Recognition sites** |
|---|---|---|
| **C50R** | for 5'- ATTTACTGG**AGG**ACAGAAGTCATCCCTCTGG -3'<br>rev 5'- GACTTCTGTCCTCCAGTAAATGGCCATGAGGATG -3' | *PshAI,*<br>introduced |
| **T142M** | for 5'- TAACA**T**GGCAAC**T**ACGGCCATGATGGTGCCCATC -3'<br>rev 5'- CGTAGTTGCCATGTTACTGATCCACATGGACAG -3' | *XcmI*, removed |
| **G219R** | for 5'- CCAGCATC**CGC**GGCACCGCCACCCTGACCGGG -3'<br>rev 5'- GCGGTGCCGCGGATGCTGGCCGCGTAGCAGATGCAC-3' | *SacII,*<br>introduced |
| **T227M** | for 5'- GACCGG**T**A**T**GGGACCCAACGTGGTGCTCC -3'<br>rev 5'- GGGTCCCATACCGGTCAGGGTGGCGGTG -3' | *AgeI,*<br>introduced |
| **S427L** | for 5'- GAGGCCT**T**GGGGCTGTCCGTGTGGATGGG -3'<br>rev 5'- CAGCCCCAAGGCCTCGGATCCTTTAGCCAG -3' | *AvaI*, removed |
| **L488P** | for 5'- GG**TT**T**A**AA**C**CCGC**C**GTACATCATGCTGCCCTGTACCC-3'<br>rev 5'- TACGGCGGGTTTAAACCGATGGAGCGAGACATGGAG -' | *PmeI,*<br>introduced |
| **Primers for real-time quantitative PCR**. | | |
| **SLC13A5** | for 5'- AAGTCATCCCTCTGGCTGTC -3'<br>rev 5'- ATCCTCTTGTGCAGGTTCCA -3' | |
| **GAPDH** | for 5'- GTCTCCTCTGACTTCAACAGCG -3'<br>rev 5'- ACCACCCTGTTGCTGTAGCCAA -3' | |

**Supplemental Table 2 The antibodies used in immunofluorescence assay, along with the specific dilutions, supplier information, and catalog numbers.**

| **Western Blotting** | | | |
|---|---|---|---|
| **Antibody** | | **Dilution** | **Company and catalog number** |

| | | | |
|---|---|---|---|
| Mouse anti-RGS(H)4, monoclonal (primary) | | 1:1000 | Qiagen # 34650 |
| Mouse anti-*Strep*-tag II, monoclonal (primary) | | 1:1500 | Qiagen # 34850 |
| Mouse Anti-SLC13A5 monoclonal (primary) | Epitope:152-VEAILQQMEATSAAT EAGLELVDKGKAKELPGSQVIFEG PTLGQQEDQERKRLCK-206 | 1:2000 | Sigma-Aldrich # SAB1402084 |
| Rabbit Anti-SLC13A5 polyclonal (primary) | Epitope: 372-SQKPKFNFRSQTEEER KTPFYPPPLLDWKVTQEKVPW-408 | 1:1000 | Sigma-Aldrich # HPA044343 |
| Rabbit Anti-SLC13A5 polyclonal (primary) | Epitope: 156-LQQMEATSAATEAGL ELVDKGKAKELPGSQVIFEGPTLGQ QEDQERKRL-204 | 1:1000 | Invitrogen # PA5-113058 |
| Recombinant rabbit anti-Ubiquitin (linkage-specific K48), monoclonal (primary) | | 1:3000 | Abcam # ab140601 |
| Rabbit anti-LC3B, polyclonal (primary) | | 1:2000 | Novus Biological #NB100-2220 |
| Anti-mouse IgG-HRP conjugate (secondary) | | 1:5000 | Sigma-Aldrich # AP308P |
| Anti-rabbit IgG-HRP conjugate (secondary) | | 1:10000 | GE Healthcare #NA934V |
| **Antibody** | **Cellular detection** | **Dilution** | **Company and catalog number** |
| Rabbit anti-B-Catenin, monoclonal (primary) | Plasma membrane | 1:1000 | Sigma Aldrich #C2206 |
| Rabbit anti-Calnexin, monoclonal (primary) | Endoplasmic reticulum | 1:1000 | Novus Biologicals #NB100-1965SS |
| Rabbit anti-GM130 monoclonal (primary) | Golgi apparatus | 1:1000 | Cell signaling #12480 |
| Rabbit anti-Lamp1, monoclonal (primary) | Lysosome | 1:400 | Cell signaling #9091 |
| Rabbit anti- Rab9a monoclonal (primary) | Endosomes | 1:50 | Cell signaling #5118 |
| Anti-mouse Alexa Flour 647(secondary) | | 1:500 | ThermoFisher Scientific # A-21235 |
| Anti-rabbit Alexa Fluor 488 (secondary) | | 1:500 | ThermoFisher Scientific # A-11008 |

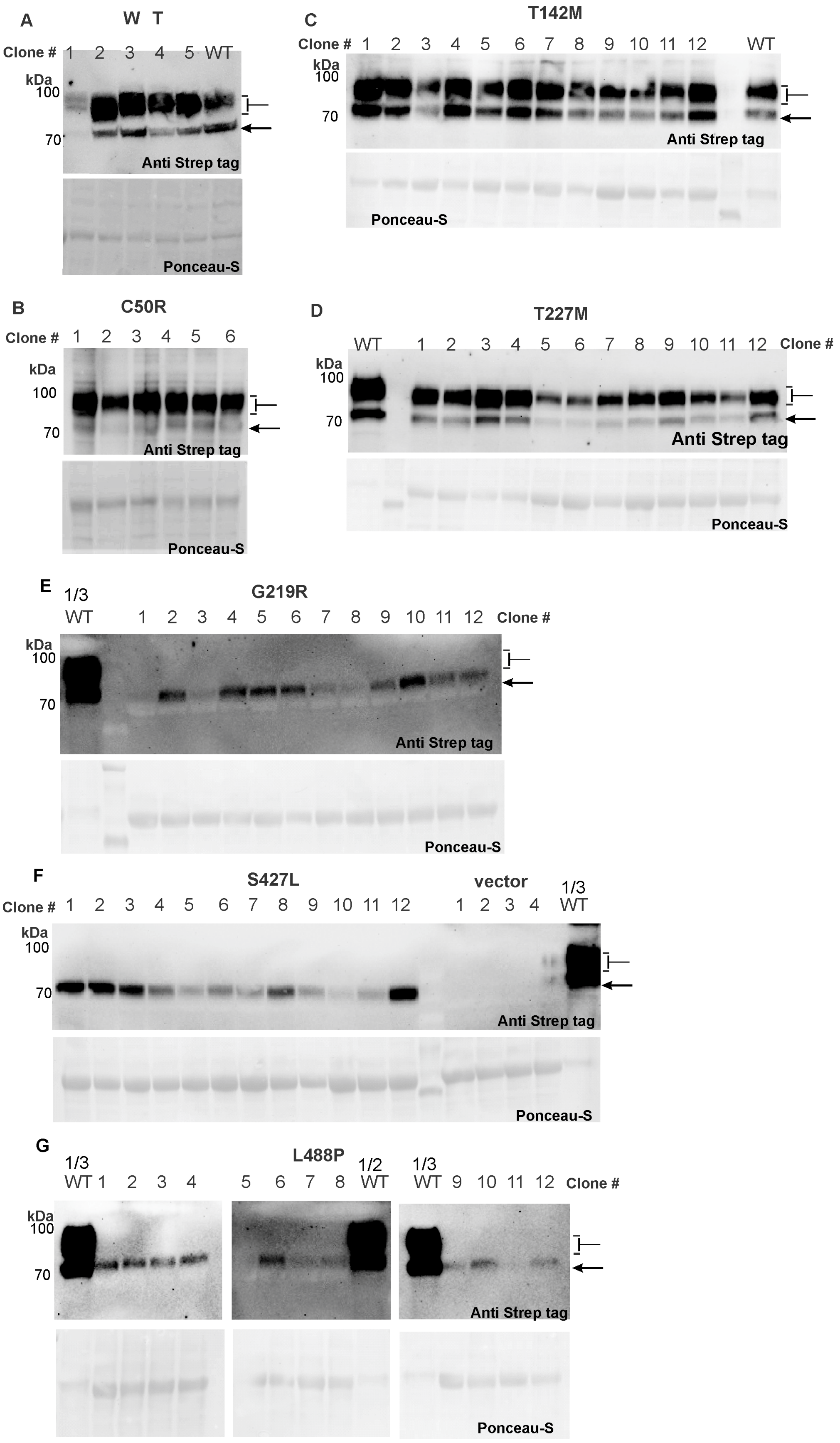

A
W T
Clone # 1 2 3 4 5 WT
kDa
100
70
Anti Strep tag
Ponceau-S
B
C50R
Clone # 1 2 3 4 5 6
kDa
100
70
Anti Strep tag
Ponceau-S
C
T142M
Clone # 1 2 3 4 5 6 7 8 9 10 11 12 WT
kDa
100
70
Anti Strep tag
Ponceau-S
D
T227M
WT 1 2 3 4 5 6 7 8 9 10 11 12 Clone #
kDa
100
70
Anti Strep tag
Ponceau-S
E
1/3 WT
G219R
1 2 3 4 5 6 7 8 9 10 11 12 Clone #
kDa
100
70
Anti Strep tag
Ponceau-S
F
S427L
Clone # 1 2 3 4 5 6 7 8 9 10 11 12
vector
1 2 3 4
1/3 WT
kDa
100
70
Anti Strep tag
Ponceau-S
G
1/3 WT 1 2 3 4
L488P
5 6 7 8 1/2 WT
1/3 WT 9 10 11 12 Clone #
kDa
100
70
Anti Strep tag
Ponceau-S

**Supplemental Figure S1 NaCT expression levels in clones selected to stably express NaCT WT, C50R, T142M, T227M, G219R, L488P, and S427L.** (**A**) Among five clonal cells transfected with WT pcDNA-HSS*-NaCT plasmid and selected for resistant to G418, four expressed higher levels of NaCT, and one expressed significantly lower levels. Clones 1 and 5 were chosen for future experiments. (**B**) Isolated C50R clones revealed five higher expressers and one lower expresser among six tested; clones 2 and 4 were selected for further experiments. (**C**) Most of the T142M clones expressed NaCT with five at high levels and seven at lower levels; clones 3 and 6 were selected for further experiments. (**D**) Similarly, most all of the T227M clones were positive with seven expressing at higher and five at lower levels; clones 5 and 3 were selected for further studies. (**E**) Among 12 clonal cells transfected with pcDNA3.1-G219R plasmid and selected with G418, five expressed higher levels and five lower levels of NaCT, while two lanes had no detectable signal. Clones 7 and 10 were selected for future experiments. (**F**) S427L showed four higher expressers and eight lower expressers; clones 6 and 12 were selected for further experiments. (**G)** L488P showed five higher expressers and five lower expressers; clones 8 and 2 were selected for further studies. Note that in (**E**), (**F**) and (**G**) only 6 μg of WT cell lysate was loaded in control lanes, one third of the total protein (20 μg) that was loaded in Class II mutant lysates. Note that all mutant lanes predominantly showed the immature, core-glycosylated NaCT protein bands at ~75 kDa (filled arrow). From the analyses, L488P expression levels appeared lowest among the three mutants analyzed.

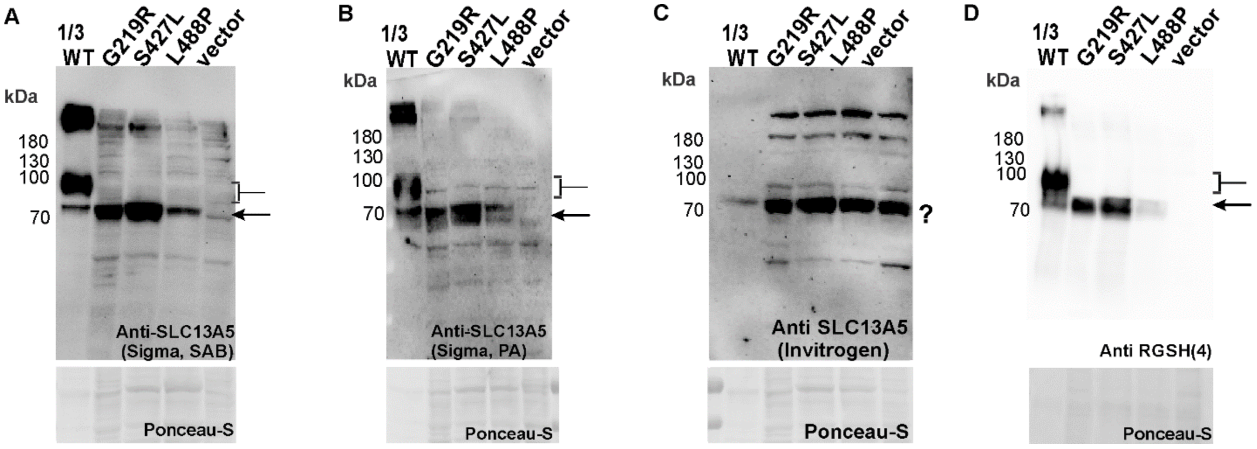


**Supplemental Figure S2 Comparison of NaCT specific antibody**. Samples of WT (clone 5), G219R (clone 10), S427L (clone 12) and L488P (clone 2) shown in Supplemntal Figure S3 were analyzed by Western blotting using commercially available Anti-SLC13A5 antibodies (**A**) SAB1402084 from Sigma Aldrich, (**B**) HPA044343 from Sigma Aldrich, and (**C**) PA5-113058 from Invitrogen; epitopes are listed in Supplemental Table S2. For comparison, the same samples were also developed with (**D**) anti-RGSH(4) specific for the engineered N-terminal epitope tag on NaCT. In principle, both anti-RGS(4) and anti Strep tag prefentially recognize the monomeric band size of the core- and complex glycosylated NaCT proteins (~75 and ~90 kDa), albeit some higher molecular weight bands >180 kDa are visible in WT lanes, which may represent oligomeric or aggregated protein. Similarly, the newly available anti-SLC13A5 antibodies (**A**) SAB1402084 and (**B**) HPA044343 from Sigma Aldrich selectively recognized the NaCT momomeric protein bands with little background observed in vector control lanes that are devoid of NaCT. In contract, antibody (**C**) PA5-113058 from Invitrogen failed to discriminate between signals in mutant and vector control lanes. Of note, Supersignal West Pico plus ECL was used to develop blots in A, C and D, while B was with Supersignal West Femto plus ECL.

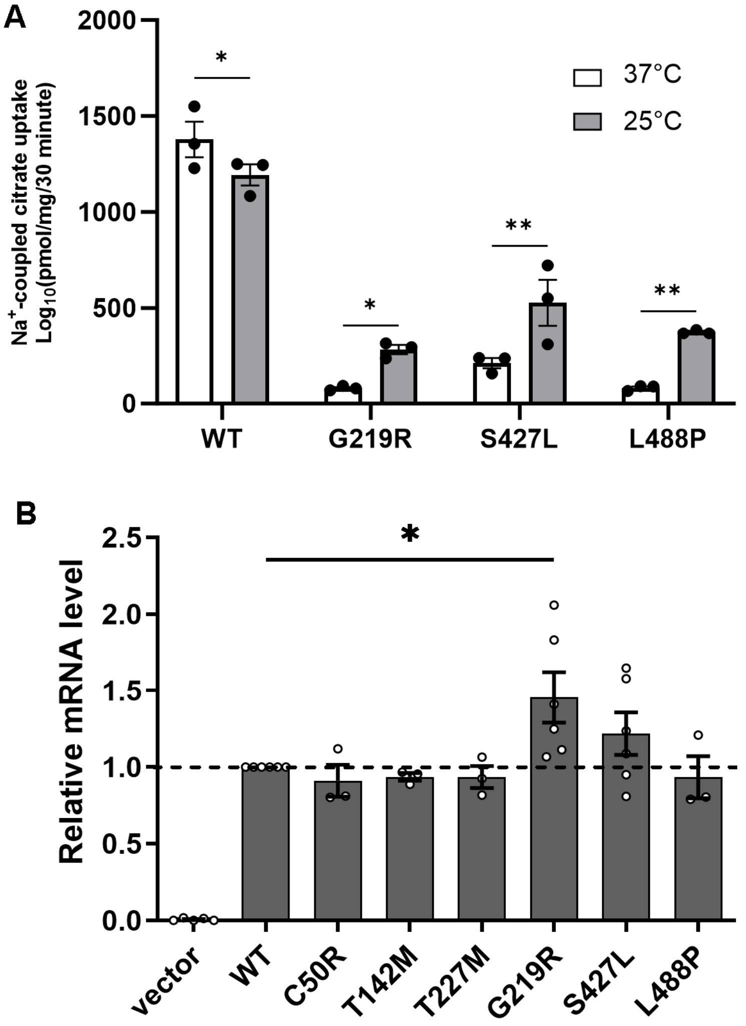


**Supplemental Figure S3 (A) Low temperature partially rescued transport activity of mutants with folding defects.** Clonal cells expressing WT or mutant proteins were cultured for 48 h days either at 37 ºC or 25 ºC, then [$^{14}$C]-citrate uptake measured for 30 at min 37 ºC in the presence of NaCl and 10 mM $Li^{+}$. Means ± SEM from three independent experiments is shown. (**B**) **mRNA levels are unchanged in NaCT mutant** cells grown at 37°C. mRNA levels were assessed by real-time quantitative PCR. No decline of mRNA was detected compared to WT, suggesting that the mutants' mRNA translation and degradation rates were unchanged. Averages ± SEM are shown from *n*= 3-12 independent experiments.

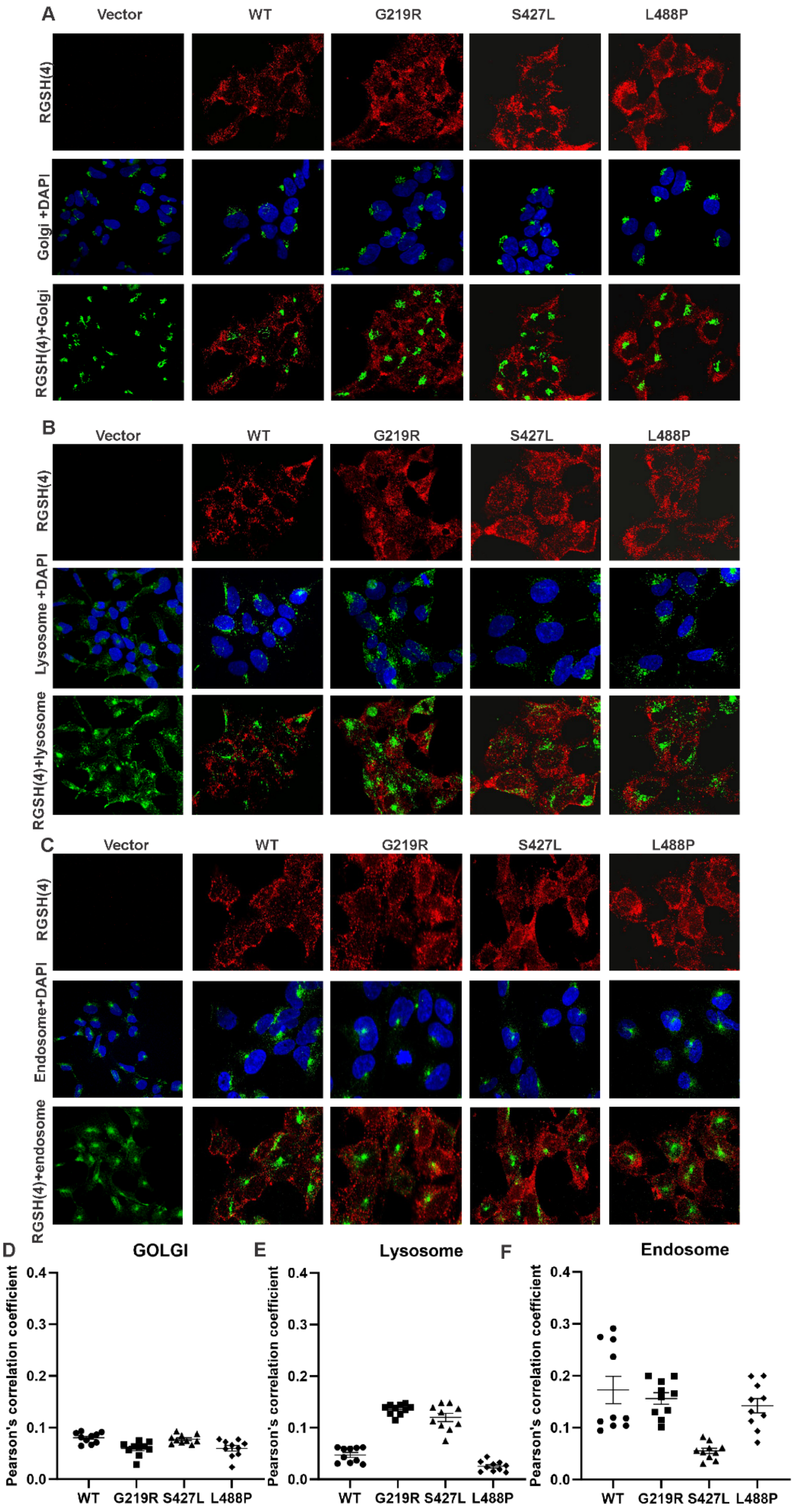

A
Vector
WT
G219R
S427L
L488P
RGSH(4)
Golgi +DAPI
RGSH(4)+Golgi
B
Vector
WT
G219R
S427L
L488P
RGSH(4)
Lysosome +DAPI
RGSH(4)+lysosome
C
Vector
WT
G219R
S427L
L488P
RGSH(4)
Endosome+DAPI
RGSH(4)+endosome
D
GOLGI
E
Lysosome
F
Endosome
Pearson's correlation coefficient
0.0
0.1
0.2
0.3
0.4
WT
G219R
S427L
L488P

**Supplemental Figure S4 Cellular localization of NaCT folding mutants using multiple cellular markers** for (**A**) Golgi apparatus, (**B**) lysosome and (**C**) endosome. (**D**), (**E**), and (**F**) are Pearson's correlation coefficient between RGSH(4) and markers. No correlation was found for any of the mutants with Golgi apparatus, lysosome or endosome markers as expected. On the other hand, some WT images showed a weak correlation with endosome marker Rab9. Means ± SEM from 10 independent images is shown.

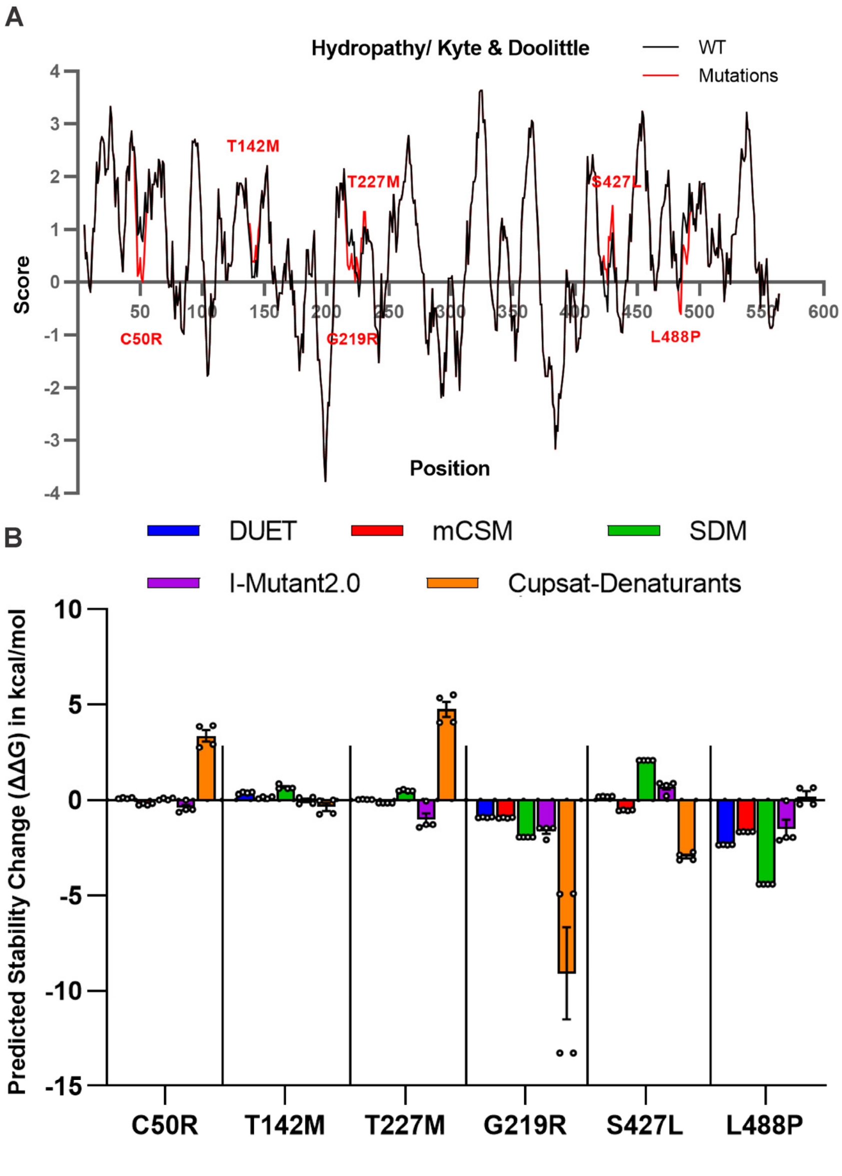


**Supplemental Figure S5 Theoretical calculations of the impacts of disease-causing mutations on hydrophobicity and protein stability.** (**A**) Kyle and Doolittle hydropathy plot. (**B**) Average of predicted stability changes (ΔΔG) in kcal/mol for C50R, T142M, G219R, T227M, S427L, and L488P computed using web servers DUET, mCSM, SDM, I-Mutant2.0 and Cupsat (denaturants choice). An important factor to understand the effects of missense mutations is their effect in protein folding and stability. However, so far there is little to no information available about NaCT missense mutations, likely because most biophysical methods to asses a protein's stability require ample source material and preferably purified mutant porteins which is an inherent challenge if protein expression and stability are compromized. Therefore, we examined available computational tools to predict mutational changes in protein stability using both chains of the two NaCT cryo-EM structures (PDB: 7JSK and 7JSJ) as input in five webservers. The in-silico protein stability of each mutation was assessed using multiple web servers, Mutation Cutoff Scanning Matrix (mCSM), Site Directed Mutator (SDM), DUET, i-Mutant2.0 and Cupsat (denaturants choice). For each mutant, input data included a PDB file (7JSK or 7JSJ), a designated chain (A or B), amino acid positions, and the specific amino acid change, resulting in four samples per mutation. Each web server utilizes a distinct approach to calculate the Predicted Stability Change ($\Delta\Delta G$) in kcal/mol. mCSM is a machine learning method that predicts missense mutation effects using structural signatures that capture distance patterns in network topology for biological systems. The SDM method uses amino acid propensities from environment-specific substitution tables for homologous protein families, providing a statistical potential energy function with an evolutionary view of the nearby residue environment. DUET offers an integrated computational approach to examine missense mutations in proteins. It combines mCSM and SDM approaches, using Support Vector Machines (SVM) to optimize and provide a consensus prediction by merging the outcomes of these distinct methods. I-Mutant2.0, a web-based tool, employs Support Vector Machine technology to predict changes in protein stability caused by single-site mutations. Utilizing data from ProTherm, a comprehensive database of experimental protein mutation information, the predictor can assess stability changes due to single-site mutations based on either protein structure or sequence. CUPSAT predicts protein stability changes due to point mutations. It utilizes amino acid-atom potentials and torsion angle distributions to analyze the mutation site's amino acid surroundings. Furthermore, it distinguishes the environment based on solvent accessibility and secondary structure specificity. Among the five methods used, Cupsat displayed significantly greater structural sensitivity. This was attributed to its energy calculation relying on statistics of torsional angles, which are highly responsive to variations in side-chain structure. Most servers predicted similar trends for a given mutation, except for S427L.
